# Transthalamic input to higher-order cortex selectively conveys state information

**DOI:** 10.1101/2023.10.08.561424

**Authors:** Garrett T. Neske, Jessica A. Cardin

## Abstract

Communication among different neocortical areas is largely thought to be mediated by long-range synaptic interactions between cortical neurons, with the thalamus providing only an initial relay of information from the sensory periphery. Higher-order thalamic nuclei receive strong synaptic inputs from the cortex and send robust projections back to other cortical areas, providing a distinct and potentially critical route for cortico-cortical communication. However, the relative contributions of corticocortical and thalamocortical inputs to higher-order cortical function remain unclear. Using imaging of cortical neurons and projection axon terminals in combination with optogenetic manipulations, we find that the higher-order visual thalamus of mice conveys a specialized stream of information to higher-order visual cortex. Whereas corticocortical projections from lower cortical areas convey robust visual information, higher-order thalamocortical projections convey strong behavioral state information. Together, these findings suggest a key role for higher-order thalamus in providing contextual signals that flexibly modulate sensory processing in higher-order cortex.

## INTRODUCTION

Many fundamental sensory, motor, and cognitive operations of the brain depend on intricate communication among multiple regions of the neocortex. Synaptic interactions among hierarchically organized neocortical regions generate increasingly selective neuronal responses ranging from simple, heterogeneous tuning properties in primary sensory areas to highly specialized responses in higher-order cortices (Orban, 2008; Riesenhuber and Poggio, 1999; Ungerleider and Mishkin, 1982). Inter-areal cortical interactions also mediate cognitive flexibility, dynamically altering the flow of information across the cortex depending on behavioral context and past experience (Haider and McCormick, 2009; Kohn et al., 2020).

Corticocortical communication comprises a complex web of feedforward and feedback connections (Felleman and Van Essen, 1991; Gămănuţ et al., 2018; Zingg et al., 2014) with distinct laminar termination patterns and cellular and subcellular targets (D’Souza et al., 2016; Harris and Shepherd, 2015; Petreanu et al., 2009). This extensive series of interactions may support the functional flexibility of cortical processing, but the relative contributions of feed-forward and feedback connections are poorly understood. Neurons in higher-level cortical areas receive robust feedforward synaptic inputs from cortical areas occupying earlier levels of the sensory processing hierarchy. However, it remains unclear to what degree higher-order sensory receptive fields arise from the integration of lower-level cortical inputs versus via selectively tuned inputs from distinct upstream subpopulations (El-Shamayleh et al., 2013; Glickfeld et al., 2013; Movshon and Newsome, 1996).

In addition to direct corticocortical innervation, higher-order thalamic projections to the cortex provide a separate pathway for corticocortical communication. Higher-order thalamic areas, including the pulvinar nucleus in the visual system, receive most of their driving synaptic input from the cortex rather than from the sensory periphery. A cortical region receiving direct, monosynaptic corticocortical connections may also receive disynaptic input from the same cortical source via higher-order thalamic nuclei (Shipp, 2003), providing a transthalamic corticocortical connection that mirrors the direct corticocortical one (Sherman and Guillery, 2011). Thalamocortical relay cells in higher-order thalamic nuclei receive very strong synaptic input from subcortically projecting layer 5 cortical pyramidal cells (Groh et al., 2008; Li et al., 2003; Reichova and Sherman, 2004), suggesting that higher-order thalamus could serve as a potent means of transmitting information between cortical regions (Theyel et al., 2010). However, the relative information carried by direct and transthalamic corticocortical connections is unclear.

Spontaneous and sensory-evoked activity in many cortical areas is strongly modulated by fluctuations in behavioral state (Benisty et al., 2023; Lohani et al., 2022; McGinley et al., 2015; Musall et al., 2019; Salkoff et al., 2020; Stringer et al., 2019; Vinck et al., 2015). Wakeful behavioral states are associated with pupil dilation, facial and body movements, locomotion, arousal, attention, and task engagement (Stringer et al 2019; Reimer et al., 2014; Vinck et al., 2015; Tang and Higley 2020). Variation in these markers of behavioral state are linked to fluctuations in cortical activity patterns and sensory encoding, as well as reorganization of cortical networks (Niell and Stryker 2010; Lohani et al., 2022; Benisty et al., 2023; Musall et al., 2023). Activity in primary thalamic relay nuclei is likewise sensitive to arousal and locomotion (Molnar et al., 2021; Spacek et al., 2022; Reinhold et al., 2023), suggesting that some behavioral signals in lower-order cortical areas may be inherited from thalamic afferents. However, partly due to the challenges of selectively manipulating activity in higher-order thalamic nuclei, it remains unclear how behavioral state information reaches higher-order cortical areas.

Here, we investigated the contributions of long-range corticocortical and higher-order thalamocortical synaptic pathways to the sensory and state-dependent properties of higher-order cortical neurons in awake, behaving mice. Using *in vivo* imaging of genetically encoded calcium indicators in neuronal cell bodies and axon terminals in layer 2/3 of higher-order mouse visual cortex, along with acute manipulations of discrete input pathways, we find that corticocortical and higher-order thalamocortical projections make distinct contributions to higher-order cortical circuit function. Whereas corticocortical inputs convey strong sensory-related information, higher-order thalamocortical inputs primarily convey contextual signals related to behavioral state. Our results provide novel insight into the synaptic foundations of corticocortical communication and suggest a unique role for higher-order thalamus in regulating flexible sensory processing by cortical circuits.

## RESULTS

### Convergence of corticocortical and higher-order thalamocortical inputs to higher visual cortex

To examine the relative contributions of cortico-cortical and thalamocortical inputs to higher-order cortical areas, we focused on the mouse posterior medial cortex (PM), a higher-order visual cortical area (Figure 1A). PM occupies one of the highest levels of the mouse visual system hierarchy (D’Souza et al., 2022; Siegle et al., 2021) and receives feedforward long-range projections from several other visual cortical regions (Wang et al., 2011 & 2012). To determine the relative abundance and density of projections to PM from different presynaptic regions, we injected a retrogradely infecting CAV2 virus (Junyent and Kremer, 2015) carrying Cre recombinase into PM of mice harboring a Cre-dependent red fluorophore. Cells in the higher-order visual thalamus, the lateral posterior nucleus (LP) (an analog of the pulvinar nucleus), but not the neighboring first-order visual thalamus, the dLGN, were robustly retro-labeled by CAV2-Cre injection in PM (Figure 1B). Projections from primary visual cortex (V1) and lateromedial cortex (LM) constituted 72–90% (mean: 82%, S.D.: 0.05%) of the feedforward corticocortical projections to PM, suggesting that projection neurons in these cortical regions may be strong drivers of PM activity. We therefore selected V1→PM and LM→PM projections for further comparison with the higher-order thalamocortical pathway from LP (i.e., LP→PM projections) (Figure 1E,F).

**Figure 1.**
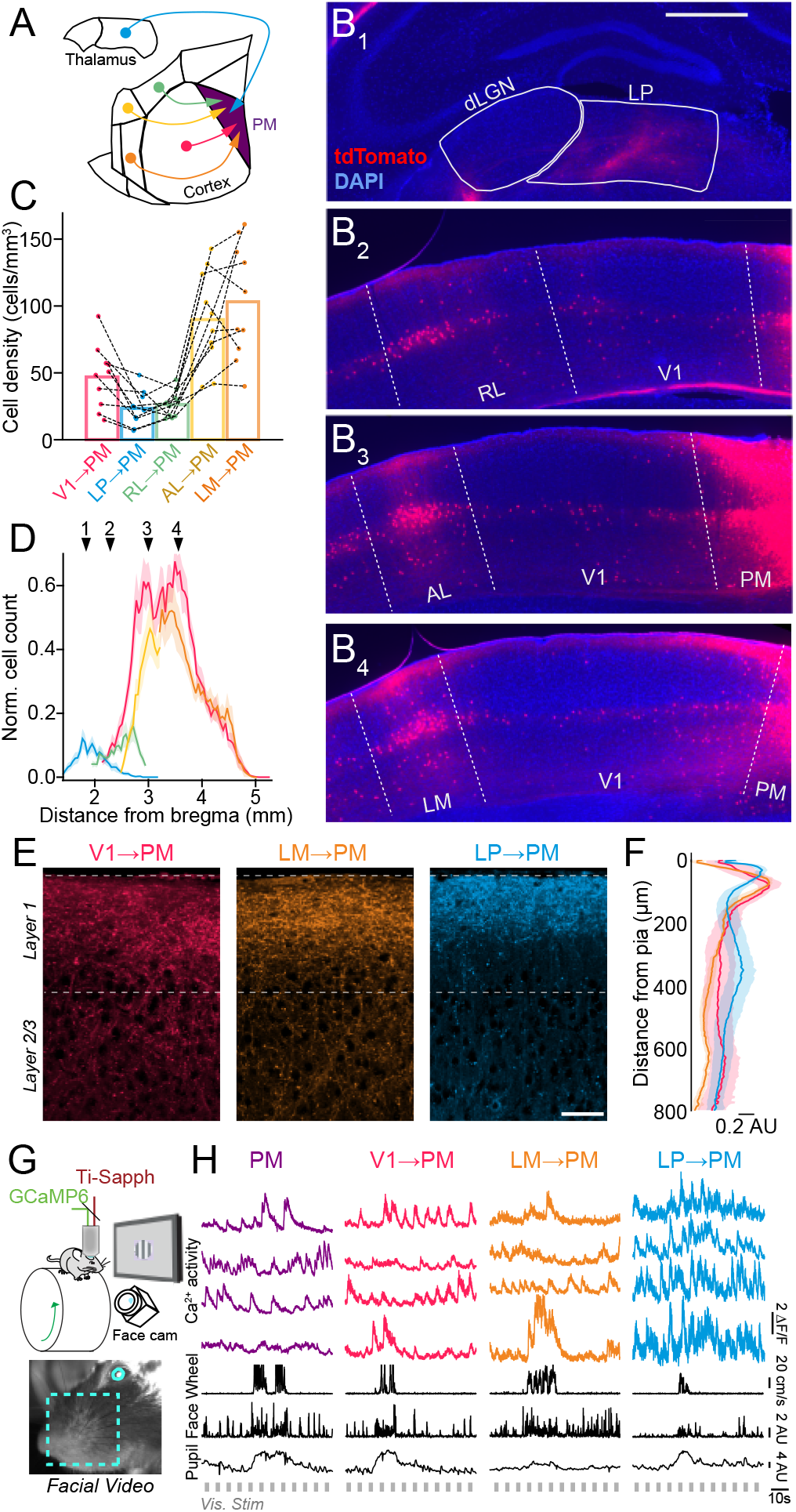
Long-range corticocortical and higher-order thalamocortical projections to mouse higher-order visual area PM. **A.** Schematic of pathways originating from the higher-order visual thalamus (lateral posterior nucleus, LP) and multiple visual cortical regions that project to the higher-order posterior medial (PM) visual cortex. **B.** Coronal sections containing thalamocortical and corticocortical projection neurons retro-labeled via injection of CAV2-Cre in PM of tdTomato reporter (Ai9) mice. Cells labeled by tdTomato send projections to PM (dLGN: dorsal lateral geniculate nucleus, RL: rostral lateral visual cortex, AL: anterior lateral visual cortex, LM: lateral medial visual cortex). Scale bar = 500 μm. **C.** Quantification of overall cell density for thalamocortical and corticocortical projection neurons sending axons to PM. Dotted lines connect values from individual Ai9 animals (N = 10 mice). **D.** Quantification of cell counts (normalized to the histological section with the highest cell count in each animal) along the anterior-posterior axis of the brain (relative to bregma) for each thalamocortical and corticocortical projection neuron type. Numbered arrows correspond to panels B_1_-B_4_ (N = 10 mice). **E.** Thalamocortical (LP→PM) and corticocortical (V1→PM, LM→PM) projection axons in PM labeled with GCaMP6s via AAV injection in the corresponding presynaptic regions. Scale bar = 50 μm. **F.** Laminar distribution of axonal fluorescence intensity for each projection type (N = 4 mice per projection type). **G.** Upper: Schematic of in vivo imaging set-up. Lower: Image from video monitoring of the mouse’s facial motion and pupil size. **H.** Fluorescence traces of Ca^2+^ activity from individual cell body (PM) and axon (V1→PM, LM→PM, LP→PM) ROIs, simultaneous with behavioral state monitoring (locomotion speed, facial motion, pupil size) and presentation of visual stimuli. Error bars denote s.e.m.

### Corticocortical and higher-order thalamocortical pathways convey distinct visual information

To directly compare the activity of cortico-cortical and thalamocortical inputs to PM with that of PM neurons, we expressed the calcium indicator GCaMP either in corticocortical and higher-order thalamocortical axons that terminate in PM or in PM neurons and assessed the state-dependent and visually evoked activity of these inputs (Figure 1 G-H, Figure S1). Compared to neurons in V1, PM neurons exhibited a nearly complete lack of tuning for visual stimulus size and a higher sensitivity to coherent motion in the visual field (Figure S2A), consistent with previous reports of enhanced motion processing in mouse dorsal visual stream areas (Sit and Goard, 2020; Wang et al., 2012).

Using a range of visual stimulation regimes (see Methods), we compared the visual response magnitudes and feature selectivity of PM cell bodies with those of V1→PM, LM→PM, and LP→PM long-range axons. Across all stimulation regimes, V1→PM axons consistently exhibited the strongest sensory-evoked responses of all the afferent populations (Figure 2). Whereas V1→PM axons and PM cell bodies were comparable in their visual response magnitudes, LM→PM and LP→PM axons exhibited significantly smaller visual response sizes for all visual stimuli (Figure 2).

**Figure 2.**
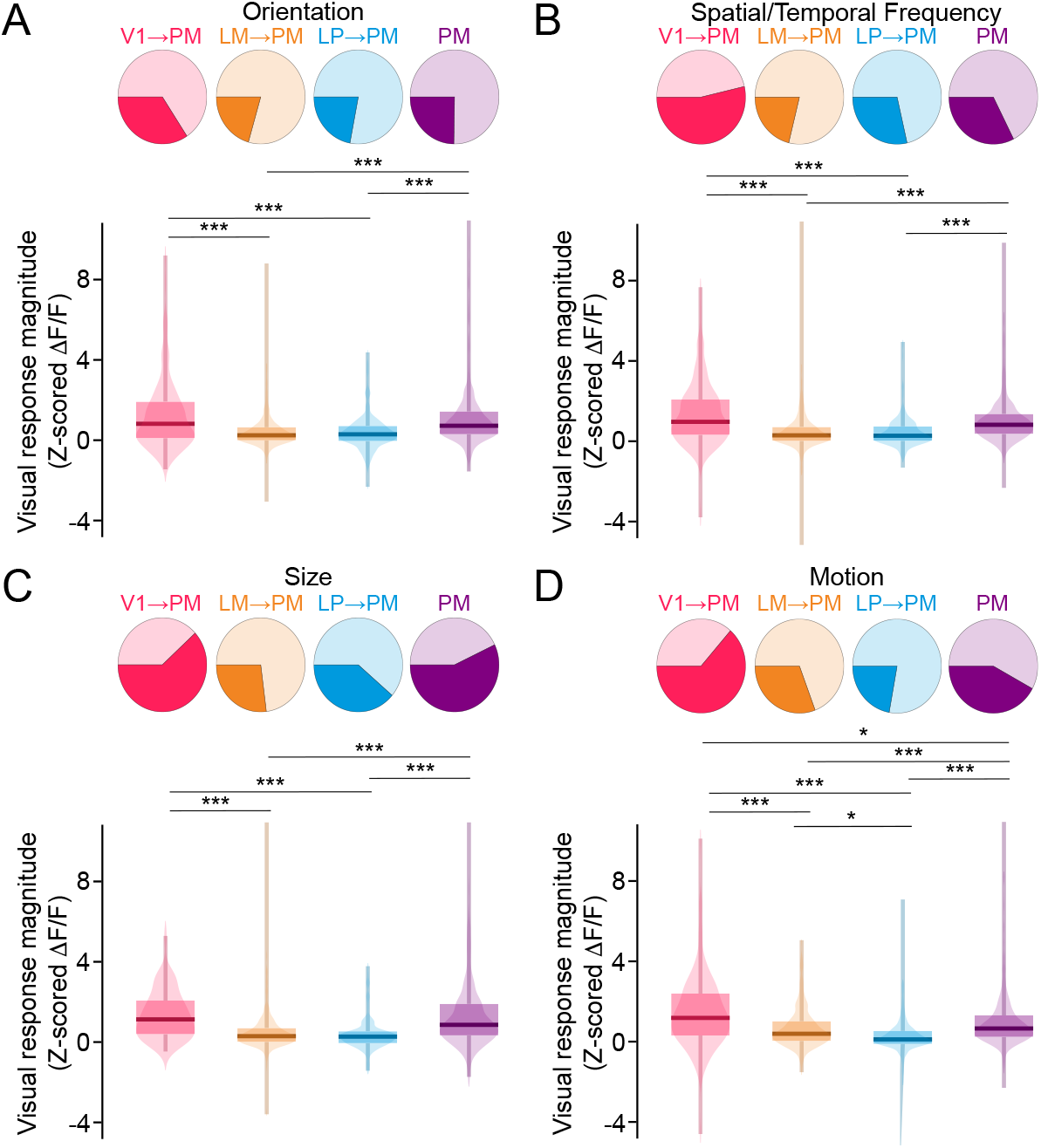
Visual responses of thalamocortical and corticocortical inputs converging in PM. **A.** Upper: ROIs of each cell type that were significantly responsive (darker colors) or non-responsive (lighter colors) to visual stimuli varying in orientation. Lower: Visual response magnitude distributions to stimuli varying in orientation for all ROIs of each cell type. (V1→PM: N = 8 animals, n = 172 ROIs; LM→PM: N = 9 animals, n = 364 ROIs; LP→PM: N = 9 animals, n = 268 ROIs; PM: N = 9 animals, n = 579 ROIs) (PM max. val. = 18.0, not shown for visualization purposes). **B.** As in A for stimuli varying in spatial frequency/temporal frequency/speed tuning. (V1→PM: N = 10 animals, n = 183 ROIs; LM→PM: N = 9 animals, n = 348 ROIs; LP→PM: N = 10 animals, n = 303 ROIs; PM: N = 13 animals, n = 781 ROIs) (LM→PM min./max. val. = −8.2/20.6). **C.** As in A but for stimuli varying in size. (V1→PM: N = 5 animals, n = 69 ROIs; LM→PM: N = 8 animals, n = 291 ROIs; LP→PM: N = 8 animals, n = 181 ROIs; PM: N = 6 animals, n = 228 ROIs) (LM→PM max. val. = 12.4; PM max. val. = 17.3). **D.** As in A but for stimuli varying in motion coherence (V1→PM: N = 7 animals, n = 200 ROIs; LM→PM: N = 7 animals, n = 301 ROIs; LP→PM: N = 10 animals, n = 376 ROIs; PM: N = 13 animals, n = 807 ROIs) (LP→PM min. val. = −30.1; PM max. val. = 12.7). *p<0.05, **p<0.01, ***p<0.001, semi-weighted t-test, Benjamini-Hochberg correction for false discovery rate.

Corticocortical and higher-order thalamocortical projections to PM exhibited diverse visual feature selectivity. For instance, whereas PM neurons largely lacked suppression in response to increasing stimulus size (Figure S2A), corticocortical and higher-order thalamocortical projections to PM exhibited robust surround suppression (Figure S2B), suggesting that the lack of suppression in PM is not inherited directly from afferents, but instead may arise via input integration (see also Murgas et al., 2020). However, the tuning properties of PM neurons were similar to those of corticocortical inputs, including V1→PM axons (Figure S2B), suggesting that some visual response features may be propagated directly from presynaptic populations in upstream visual cortical areas (Glickfeld et al., 2013).

### Higher-order thalamocortical pathways convey state-related information

The spontaneous and sensory-evoked activity of neurons across the entire cortex is strongly influenced by behavioral or arousal state. Changes in wakeful behavioral states are often associated with motor actions, such as locomotion, facial, and whisker motion, and altered pupil size (Fu et al., 2014; McGinley et al., 2015; Musall et al., 2019; Neske et al., 2019; Niell and Stryker, 2010; Polack et al., 2013; Reimer et al., 2014; Salkoff et al., 2020; Stringer et al., 2019; Vinck et al., 2015). Although ascending neuromodulatory systems play a crucial role (Lohani et al., 2022), both corticocortical and thalamocortical afferents could also be key players in cortical state modulation (Nestvogel and McCormick, 2022; Poulet et al., 2012; Zagha et al., 2013). We therefore considered what roles the major feedforward corticocortical (V1→PM and LM→PM) and higher-order thalamocortical (LP→PM) inputs may play in conveying state-dependent activity to PM.

PM neurons exhibited stronger state-dependent modulation than V1 neurons for locomotion (Figure S3A,B), and for facial motion (Figure S3C-E), and pupil diameter (Figure S3F-H) in the absence of locomotion. Consistent with this observation, V1→PM axons exhibited the weakest levels of overall state-dependent modulation (Figure 3B-I), in direct contrast with their robust modulation by visual input. State-dependent modulation was stronger for higher-order thalamocortical inputs from LP (LP→PM axons) than either of the corticocortical afferents (V1→PM and LM→PM axons) (Figure 3B-I). Together, these results suggest that whereas corticocortical axons from V1 carry the strongest sensory-related information to PM, higher-order thalamocortical axons from LP carry the strongest state-related information.

**Figure 3.**
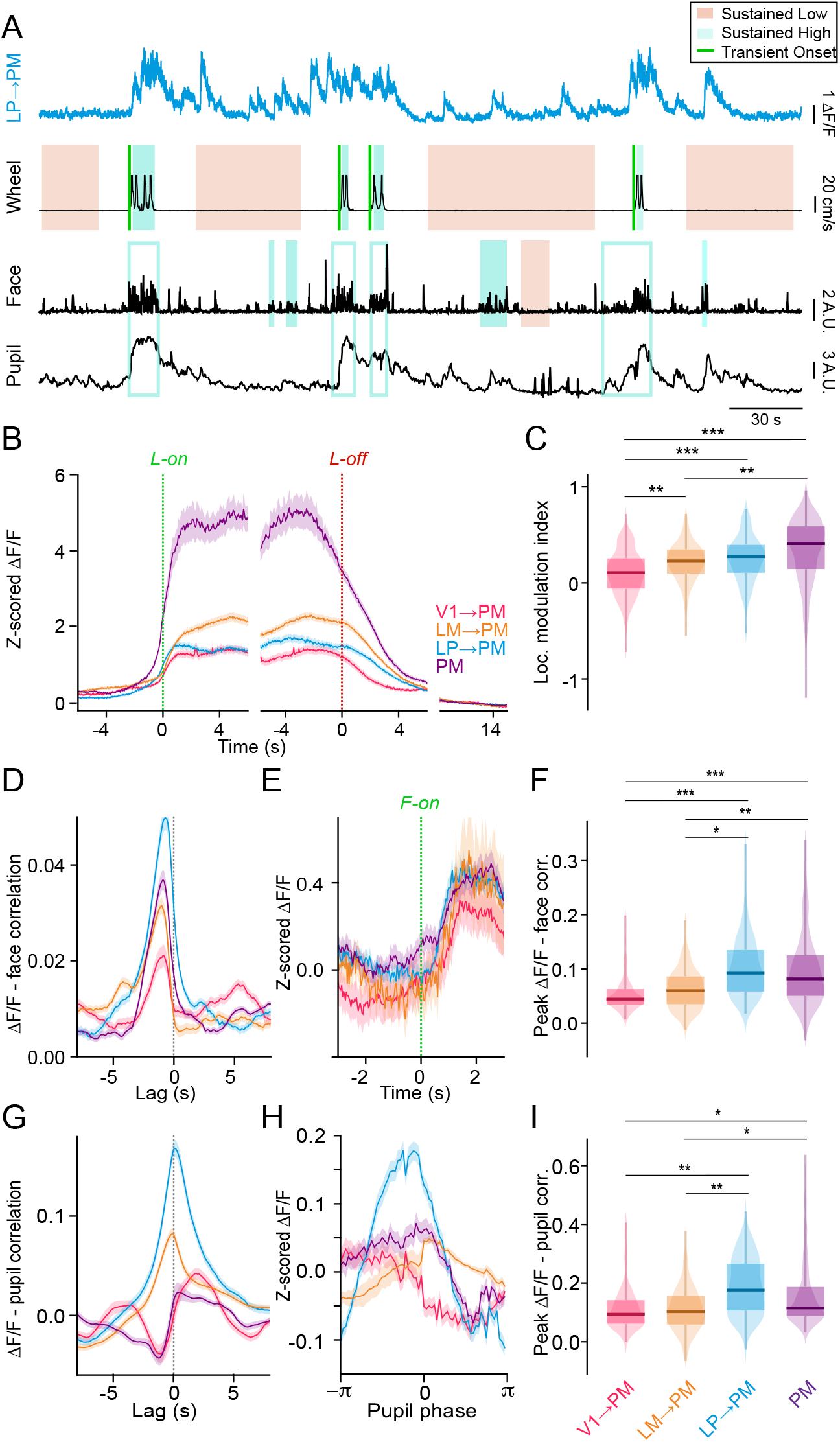
Behavioral state-dependent modulation of thalamocortical and corticocortical inputs to PM. **A.** Example Ca^2+^ signal from an LP axon terminating in PM (cyan). Lower traces (black) show locomotion speed, facial motion (motion energy of the whisker pad), and pupil size (AU, arbitrary units). Periods of sustained low or high motor activity (locomotion or facial motion) are indicated by shaded areas and transition points from low to high motor activity are indicated by green lines. Analyses of modulation by facial motion and pupil dilation/constriction were limited to sustained periods without locomotion. Open blue boxes indicate high facial motion periods not included in the analysis of state transitions due to overlap with locomotion. **B.** Neuronal activity (Z-scored ΔF/F) aligned to locomotion onset (green dotted line) and offset (red dotted line). Baseline activity (10-15 s after locomotion offset during sustained quiescent periods) is shown to the right for comparison. **C.** Locomotion modulation indices for ROIs of each cell type (V1→PM: N = 11 animals, n = 291 ROIs; LM→PM: N = 9 animals, n = 343 ROIs; LP→PM: N = 10 animals, n = 266 ROIs; PM: N = 8 animals, n = 622 ROIs). **D.** Cross-correlation between neuronal activity (ΔF/F) and facial motion. Facial motion is the reference signal in the cross-correlation. **E.** Neuronal activity (Z-scored ΔF/F) aligned to the onset of high facial motion. **F.** Peak correlation values between neuronal activity and facial motion for each cell type (V1→PM: N = 4 animals, n = 107 ROIs; LM→PM: N = 4 animals, n = 176 ROIs; LP→PM: N = 6 animals, n = 165 ROIs; PM: N = 5 animals, n = 454 ROIs). **G.** Cross-correlation between neuronal activity (ΔF/F) and pupil size. **H.** Neuronal activity (Z-scored ΔF/F) aligned to one pupil dilation-constriction cycle (derived from the Hilbert transform of the pupil signal). **I.** Peak correlation values between neuronal activity and pupil size for each cell type (V1→PM: N = 4 animals, n = 108 ROIs; LM→PM: N = 8 animals, n = 284 ROIs; LP→PM: N = 6 animals, n = 162 ROIs; PM: N = 5 animals, n = 353 ROIs). *p<0.05, **p<0.01, ***p<0.001, semi-weighted t-test, Benjamini-Hochberg correction for false discovery rate. Error bars denote s.e.m.

The higher-order thalamic nucleus LP receives convergent synaptic inputs from multiple cortical and subcortical brain regions (Blot et al., 2021), as well as from multiple neuromodulatory systems (McCormick, 1992; Varela, 2014). Previous anatomical work has shown that synaptic input to PM-projecting LP neurons largely arises from corticothalamic neurons in V1 (Blot et al., 2021). We therefore compared the strength of the state-dependent modulation between two classes of V1 projection neurons: 1) corticocortical V1 neurons projecting directly to PM and 2) corticothalamic V1 neurons projecting to LP (Figure S4A,B). The somata of these pyramidal neurons largely resided in layer 5 and lower layer 6 (Figure S4C,D). As a proxy for calcium activity in the deep-layer somata, we imaged the calcium activity in the layer-1 apical dendrites of these neurons (Beaulieu-Laroche et al., 2019; Peters et al., 2017) (Figure S4E,F). Compared to V1 corticocortical projection neurons, V1 corticothalamic neurons projecting to LP exhibited stronger modulation by behavioral state, specifically for state transitions associated with dilations in pupil diameter during behavioral quiescence (Figure S4L-N). V1 projection neurons may thus be a key source of the state-dependent activity observed in LP→PM projections.

### Functional interactions between afferents and PM neurons

To examine the functional contributions of corticocortical and thalamocortical inputs to PM, we simultaneously imaged the calcium activity of PM cell bodies and the axon terminals of the V1→PM, LM→PM, or LP→PM projections. To separate axons from local neuropil, we expressed GCaMP6s in axon terminals and a soma-targeted version of GCaMP, riboGCaMP6m (Chen et al., 2020), in PM cell bodies (Figure 4A,B). We observed very little GCaMP labeling in the axons or dendrites of PM neurons when riboGCaMP6m was expressed in these cells alone (Figure S5A). Furthermore, imaging of *in vivo* preparations in which only riboGCaMP6m was expressed in PM cell bodies showed negligible neuropil signal when compared with GCaMP6s (Figure S5B-E). This dual GCaMP approach allowed simultaneous imaging of cell bodies and axons from a single imaging channel (Figure 4A-C).

**Figure 4.**
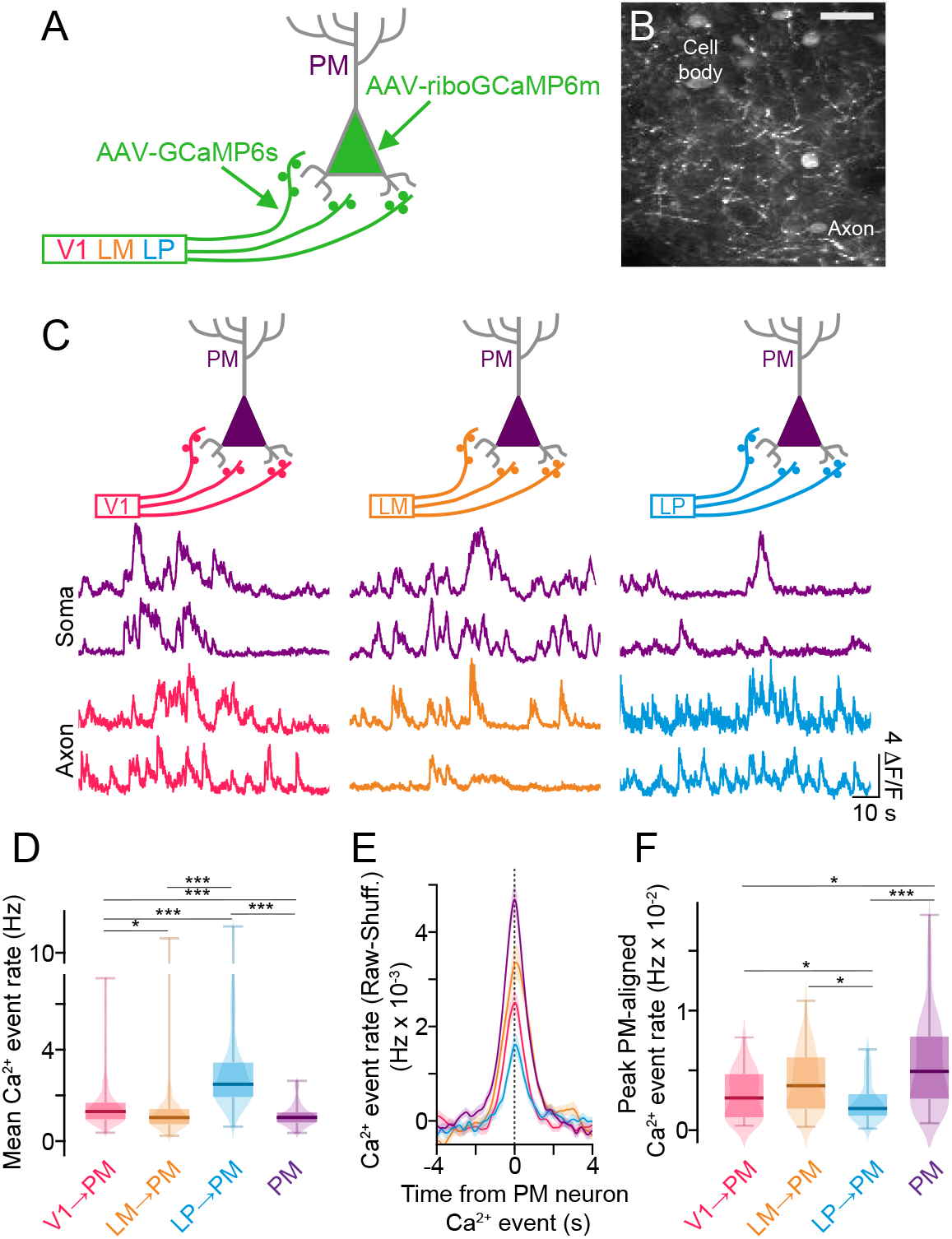
Distinct functional interactions between thalamocortical and corticocortical afferents and PM cells. **A.** Schematic of method for simultaneously monitoring the neuronal activity of PM cell bodies and long-range projection axons terminating in PM, based on expression of GCaMP6s in long-range axon terminals and ribo-GCaMP6m in PM cell bodies. **B.** In vivo field of view with PM cell bodies expressing riboGCaMP6m and LM axon terminals expressing GCaMP6s. **C.** Fluorescence traces of Ca^2+^ activity simultaneously recorded in PM cell bodies expressing riboGCaMP6m and projection axons expressing GCaMP6s. **D.** Mean Ca^2+^ event rates among the different cell types. (V1→PM: N = 5 animals, n = 332 ROIs; LM→PM: N = 4 animals, n = 211 ROIs; LP→PM: N = 5 animals, n = 191 ROIs; PM: N = 14 animals, n = 168 ROIs). **E.** Ca^2+^ event rates for each afferent population aligned to simultaneously recorded Ca^2+^ events in PM neurons (raw data minus data from shuffled PM event times). **F.** Peak Ca^2+^ event rates aligned to PM neuron Ca^2+^ events. (V1→PM: N = 5 animals, n = 59 PM ROIs; LM→PM: N = 4 animals, n = 53 PM ROIs; LP→PM: N = 5 animals, n = 56 PM ROIs; PM activity aligned with PM events: N = 14 animals, n = 168 PM ROIs). *p<0.05, **p<0.01, ***p<0.001, semi-weighted t-test, Benjamini-Hochberg correction for false discovery rate. Error bars denote s.e.m.

Using this approach, we examined the relationship between neuronal activity in PM cell bodies and V1→PM, LM→PM, and LP→PM axons during quiescent (non-locomotion) periods. After deconvolving ΔF/F calcium activity traces from axons and cell bodies into estimates of calcium events (Materials and Methods), we assessed the relationship between simultaneously imaged axonal and PM somatic calcium event rates. Although overall calcium event rates in LP axons were significantly higher than those of V1 or LM corticocortical axons and PM cell bodies (Figure 4D), LP events were significantly less time-locked to events in PM cell bodies than those of cortico-cortical axons (Figure 4E,F). LP→PM activity may thus be less effective in driving PM neuron spiking than corticocortical inputs. Indeed, functional interactions between LP→PM axons and PM cell bodies were comparable with those between LP→V1 axons and V1 cell bodies, a canonical modulatory pathway (Figure S6) (Miller-Hansen and Sherman, 2022; Roth et al., 2016). Calcium events in PM cell bodies were significantly more correlated with events in neighboring PM cell bodies than to those in either corticocortical or thalamocortical axons (Figure 4E,F), potentially reflecting strong levels of recurrent connectivity within PM (Li et al., 2021). Together, these results support a modulatory role for LP inputs to PM and a stronger driving role for short-and long-range corticocortical inputs.

### Acute silencing reveals distinct causal contributions to PM neuronal activity

To determine more precisely the contributions of each input to the activity of PM neurons, we selectively silenced individual inputs to PM while monitoring the state-dependent and visually evoked activity of PM neurons. Using cre-dependent expression of the inhibitory opsin eOPN3 (Mahn et al., 2021), we silenced the axon terminals of different corticocortical and higher-order thalamocortical projections within PM. To express eOPN3 selectively in V1→PM, LM→PM, or LP→PM terminals, we injected a mixture of CAV2-Cre and AAV-GCaMP6s into PM followed by the Cre-dependent AAV-SIO-eOPN3-mScarlet in the presynaptic target area (Figure 5A). These injections resulted in projection-specific expression of eOPN3 in axon terminals that were physically intermingled with PM cell bodies expressing GCaMP6s (Figure 5B). The relative of numbers of V1→PM, LM→PM, or LP→PM projection neurons labeled in this way largely recapitulated those seen from CAV2-Cre injection in reporter animals (Figure S7C), indicating that eOPN3 was not over- or under-expressed in any projection neuron class. Visual responses of PM neurons were robustly decreased by optogenetic suppression of corticocortical V1 and LM axons, but only moderately affected by suppression of thalamocortical LP axons (Figure 5C,D,G). Consistent with the strong state-dependent modulation of LP axons we observed in PM, state-dependent modulation of PM neuron activity with changes in pupil diameter was significantly reduced when LP→PM, but not V1→PM or LM→PM, terminals were inhibited (Figure 5E,F,H) (Figure S7D-I). Corticocortical and higher-order thalamocortical inputs may thus confer the sensory- and state-dependent properties of PM cortical neurons, respectively.

**Figure 5.**
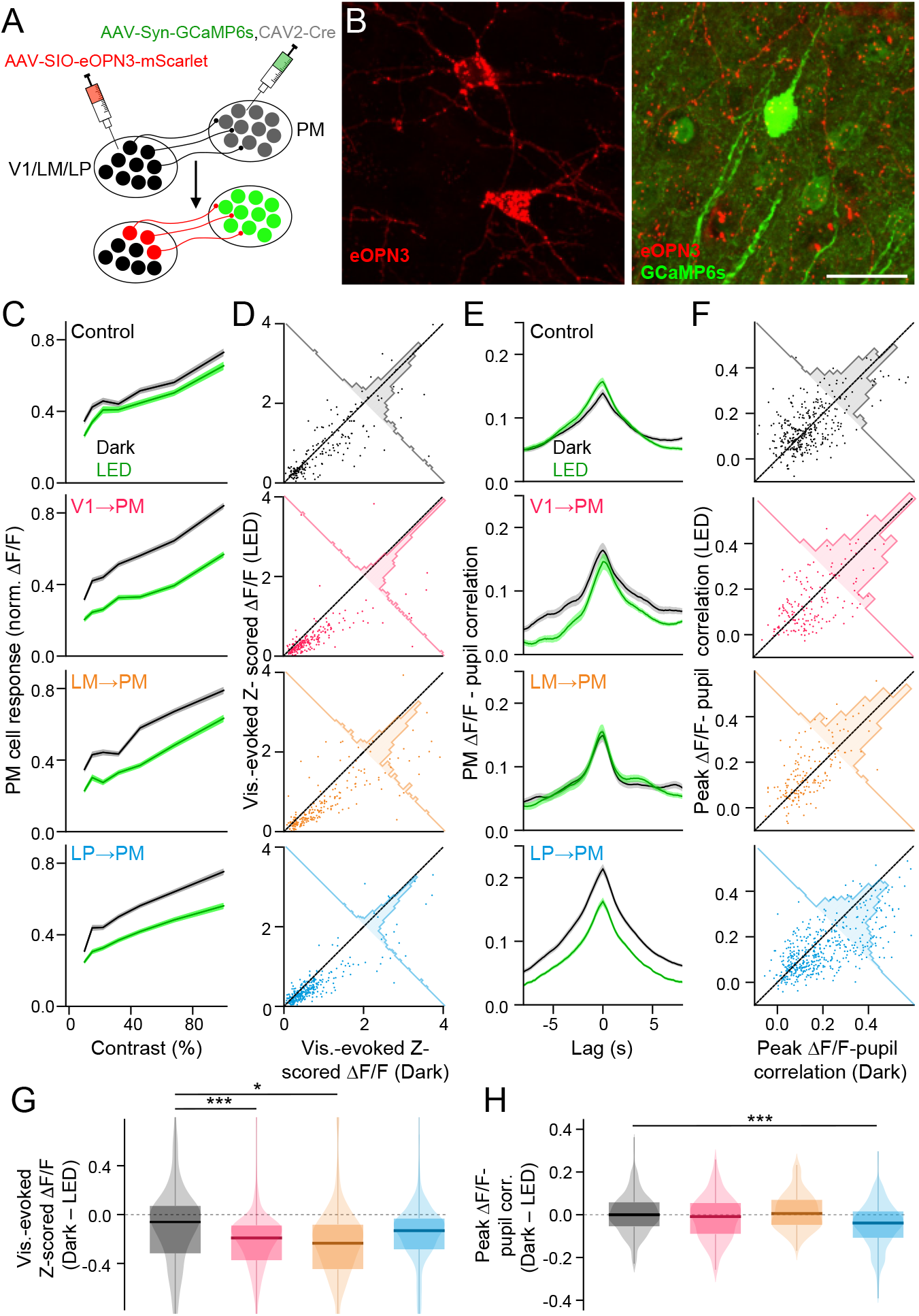
Distinct contributions of corticocortical and thalamocortical pathways to PM activity. **A.** Schematic of viral injections for co-expressing GCaMP6s in PM cortical neurons and the Cre-dependent inhibitory opsin eOPN3 in presynaptic projection axon terminals. **B.** Left: Expression of eOPN3-mScarlet (red) in thalamocortical neurons in LP. Right: Expression of GCaMP6s in PM cortical neurons (green) and eOPN3-mScarlet (red) in the surrounding LP thalamocortical terminals. Scale bar = 30 μm. **C.** Visual contrast-response curves of PM neurons in control animals and animals with eOPN3 expressed in corticocortical and higher-order thalamocortical axons for session with (LED) and without (Dark) optogenetic stimulation. **D.** Scatter plots showing visual response magnitudes from individual PM neurons identified during both Dark and LED imaging sessions. **E.** As in C but for cross-correlations between PM neuronal activity and pupil size. **F.** As in D but for peak correlation values between neuronal activity and pupil size. **G.** Differences between visual response magnitudes during sessions with and without optogenetic stimulation. (Control: N = 5 animals, n = 195 ROIs; V1→PM eOPN3: N = 7 animals, n = 227 ROIs; LM→PM eOPN3: N = 4 animals, n = 215 ROIs; LP→PM eOPN3: N = 6 animals, n = 327 ROIs) (Control min./max. val. = −3.2/2.5; V1→PM eOPN3 min./max. val. = −5.3/3.1; LM→PM eOPN3 min./max. val. = −3.1/5.0; LP→PM eOPN3 min./max. val. = −3.1/1.0). **H.** As in G but for differences in peak ΔF/F-pupil correlation values between sessions with and without optogenetic stimulation. (Control: N = 7 animals, n = 327 ROIs; V1→PM eOPN3: N = 4 animals, n = 169 ROIs; LM→PM eOPN3: N = 3 animals, n = 167 ROIs; LP→PM eOPN3: N = 5 animals, n = 468 ROIs). *p<0.05, **p<0.01, ***p<0.001, semi-weighted t-test, Benjamini-Hochberg correction for false discovery rate. Error bars denote s.e.m.

**Figure 6.**
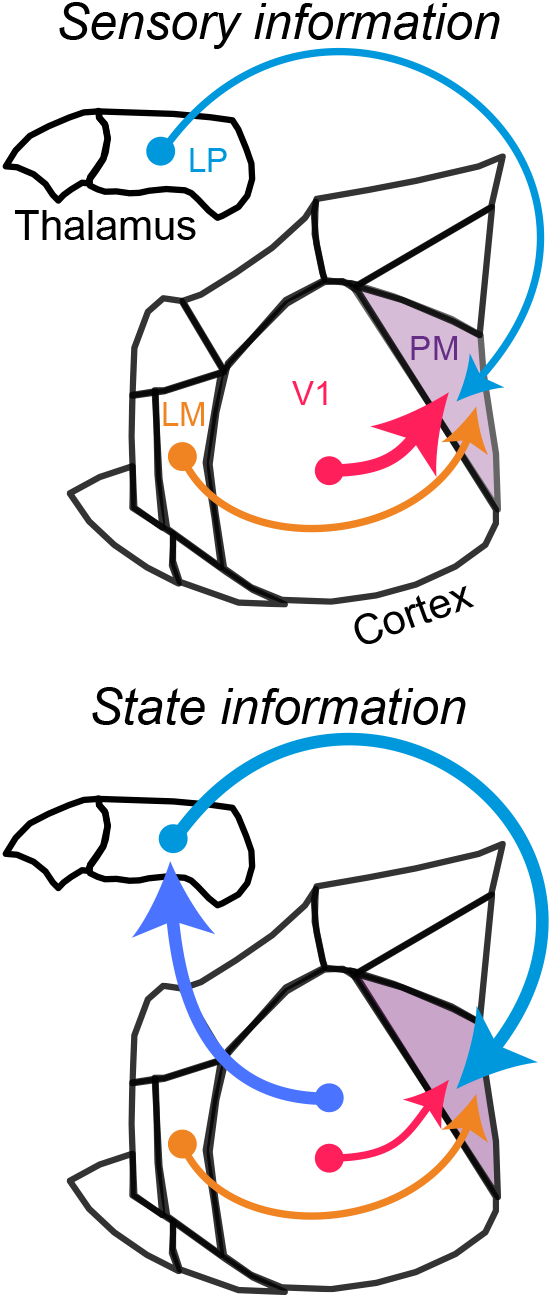
Corticocortical and thalamocortical pathways provide distinct streams of information to higher-order cortex. Upper: Corticocortical pathways, particularly the direct input from V1, carry strong sensory information to higher-order visual cortex (PM), whereas higher-order thalamocortical inputs from LP carry comparatively weak sensory signals. Lower: Corticocortical pathways carry relatively little behavioral state information, whereas higher-order thalamocortical inputs from LP convey strong behavioral state-related signals that reflect their presynaptic corticothalamic inputs from V1.

## DISCUSSION

Our results reveal distinct roles for corticocortical and higher-order thalamocortical pathways to higher-order cortex. We examined the sensory responses and behavioral state-dependent modulation of activity in two lower-order visual cortical areas, V1 and LM, and a higher-order thalamocortical area, LP, that converge on a common higher-order visual cortical area, PM. We find that while corticocortical projections carry strong sensory information and significantly impact the visually-evoked responses of their postsynaptic targets in PM, higher-order thalamocortical signals from LP do not strongly contribute to PM visual responses. In contrast, direct corticocortical inputs from V1 and LM do not drive state-dependent modulation in PM, whereas inputs from LP contribute to the representation of behavioral state by PM neurons. Together, these results suggest two distinct streams conveying sensory and contextual information to higher-order cortex.

Unlike first-order thalamic nuclei, which receive their driving synaptic input from the sensory periphery or associated brainstem nuclei, most higher-order thalamic nuclei receive their strongest input from the cortex. Anatomical connectivity suggests that whereas first-order thalamic nuclei (e.g., dLGN) predominately function as sensory relays, albeit under dynamic modulatory control from the cortex (Crandall et al., 2015; Spacek et al., 2022; Reinhold et al., 2023), higher-order thalamic nuclei (e.g. pulvinar) relay the results of computations carried out within local cortical networks. Like first-order thalamus, which encodes low-level sensory information from the sensory periphery, higher-order thalamic nuclei may encode integrated sensory information that is propagated across the cortex (Merabet et al., 1998). Alternatively, higher-order thalamic nuclei may predominantly convey non-sensory information or reconfigure the cortical circuits to which they project (Schmitt et al., 2017). Previous work has suggested that the broad tuning properties of pulvinar outputs seem inadequately suited to generate the highly specific receptive fields of extrastriate neurons (Van Essen, 2005), but may instead play an essential role in modifying the functional connectivity of distributed cortical regions in a task-dependent manner (Fiebelkorn et al., 2019; Saalmann et al., 2012; Zhou et al., 2016).

Targeted imaging of corticortical inputs to PM and of PM neurons revealed robust visual responses (Andermann et al., 2011; Murgas et al., 2020; Blot et al., 2021). Overall, PM neurons exhibited low selectivity for stimulus orientation, enhanced motion sensitivity, and almost no surround suppression as compared to V1 neurons. In good agreement with previous reports (Bennett et al., 2019; Blot et al., 2021), we observed that LP afferents likewise showed visual responses, but with low selectivity for stimulus orientation and little size tuning. LP afferents to PM exhibited high levels of activity but little correlation with the activity of individual PM neurons, consistent with a modulatory rather than driving role for higher-order thalamic inputs.

Surprisingly, we found that PM neurons were more strongly modulated by changes in behavioral state, as measured by locomotion, facial motion, and pupil dilation, than the general population of V1 neurons or any individual direct afferent pathway. This strong representation of behavioral state by PM neurons may facilitate non-visual roles of PM in visually guided tasks (Jin and Glickfeld, 2020; Goltstein et al., 2021). LP inputs to PM exhibited more robust sensitivity to behavioral state variables than either V1 or LM inputs, suggesting that higher-order thalamus may provide a key source of state information to PM. Indeed, LP thalamocortical activity was robustly linked to pupil-linked arousal and optogenetic inactivation of LP terminals reduced the modulation of postsynaptic PM neurons by pupil dilation. Consistent with a contextual role for LP, recent work found that LP thalamocortical axons projecting to higher visual cortex do not encode a detailed representation of optic flow, which is signaled by corticocortical axons, but rather discrepancies between optic flow and self-motion (Blot et al., 2021). Corticocortical afferents may thus be a principal conduit of feedforward sensory information between cortical areas, whereas higher-order thalamocortical axons may primarily contribute state-dependent contextual modulation of this sensory signaling (Petty et al., 2021; La Terra et al., 2022).

Deep-layer cortical projection neurons may be a major source of the state-dependent modulation of LP activity. Although V1 projects to both PM and LP, these pathways arise from distinct populations of projection neurons with distinct properties (Lur et al., 2016; Tang and Higley 2020). We found that V1 layer 5 corticothalamic neurons that project to LP are more strongly modulated by behavioral state than corticocortical projection neurons. Previous work has likewise highlighted a potential role for corticothalamic projections in conveying locomotion modulation signals from V1 to LGN (Reinhold et al., 2023). Additional pre-synaptic sources may further amplify the unique state-dependent properties of LP thalamocortical neurons. Previous work has identified several neuromodulatory systems (Varela and Sherman, 2007, 2009) and GABAergic pathways (Bokor et al., 2005; Martinez-Garcia et al., 2020) that selectively interact with higher-order thalamocortical neurons. State information may also reach LP via corticothalamic neurons in the deepest part of cortical layer 6, which project exclusively to higher-order thalamus (Hoerder-Suabedissen et al., 2018) (see also Figure S4C,D). Deep layer 6 corticothalamic neurons are sensitive to the wake-promoting neuropeptide orexin (Bayer et al., 2004), which is strongly coupled to pupil-linked arousal (Grujic et al., 2023). Neuromodulatory signaling to corticothalamic neurons may thus regulate a selective channel for behavioral state information from V1 to LP, potentially contributing to the strong correlation between pupil fluctuations and LP activity.

Our current results pertain to long-range synaptic inputs to a specific region of the higher-order mouse visual cortex (area PM). However, the mouse visual system consists of more than 10 distinct cortical processing regions downstream of V1 (Zhuang et al., 2017). These regions have specific hierarchical relationships and communicate in both feedforward and feedback directions (D’Souza et al., 2022; Glickfeld and Olsen, 2017; Wang and Burkhalter, 2007; Wang et al., 2011). Furthermore, analogously to the primate pulvinar (Kaas and Lyon, 2007), mouse LP contains multiple retinotopically organized subdivisions (Bennett et al., 2019). The functional role of LP in corticocortical communication may therefore differ depending on the hierarchical positions of the postsynaptic cortical regions. For instance, whereas LP input to V1 (Roth et al., 2016) and medial higher visual areas (such as PM) may primarily convey contextual and behavioral state signals, LP input to lateral higher visual areas (such as the postrhinal cortex) may instead convey robust and precise visual information (Beltramo and Scanziani, 2019). LP thalamocortical neurons might also have distinct functions based upon which specific pre-and postsynaptic cortical areas they connect, i.e., which transthalamic pathway they complete. Furthermore, the synaptic mechanisms by which higher-order thalamocortical inputs from transthalamic pathways are integrated with direct corticocortical inputs within the cortex, and how this integration contributes to cortical circuit function, remain to be explored.

Previous work has shown that activity in ascending neuromodulatory pathways, such as those releasing acetylcholine and norepinephrine, may be linked to pupil-related arousal (Reimer et al., 2016). Neuromodulatory pathways may also carry motor-related signals, such as locomotion and facial motion (Collins et al., 2023; Lohani et al., 2022). Our current results indicate that higher-order thalamocortical axons convey state-modulated signals reminiscent of canonical neuromodulatory pathways. Due to their more restricted axonal arborization and ionotropic glutamatergic signaling, higher-order thalamocortical projections might provide a more spatiotemporally precise form of state modulation compared to cholinergic and noradrenergic pathways. Future work will require disentangling the precise contributions of these different afferent pathways to both global and local state-dependent reconfigurations of cortical networks.

Overall, our findings identify corticocortical and higher-order thalamocortical channels that convey distinct information streams to higher-order cortex. We find that corticocortical pathways convey robust sensory information, whereas higher-order thalamocortical pathways primarily provide behavioral state information. Higher-order thalamocortical inputs may thus serve as a selective route for contextual information to shape the dynamic functional interactions between cortical regions and the integration of sensory information to guide behavioral output.

## MATERIALS AND METHODS

### Animals

All animal handling and maintenance was performed according to the regulations of the Institutional Animal Care and Use Committee of the Yale University School of Medicine. Adult male and female C57BL/6J mice (The Jackson Laboratory) aged 10–14 weeks were kept on a 12h light/dark cycle, provided with food and water ad libitum, and housed individually following headpost implants. All experiments were performed during the light cycle.

### Surgical procedures

Surgeries were performed in adult mice (P90–P180) in a stereotaxic frame, anesthetized with 1-2% isoflurane mixed with pure oxygen. Eyes were protected throughout the surgery with ophthalmic ointment (Puralube). After providing an intraperitoneal injection of Carprofen (5 mg/kg), a midline incision along the scalp was made to expose the skull. Small burr holes in the skull were made in stereotaxic coordinates to introduce viral vectors into multiple brain regions of interest– (mm lateral [L], posterior [P], and ventral [V] from bregma) LP thalamus: 1.6 L, 2.06 P, 2.7 V; PM visual cortex (2 sites): 1.8L, 2.7P, 0.4V & 1.5L, 2.9P, 0.4V; V1 visual cortex (3 sites): 2.5L, 3.5P, 0.4V & 3.0L, 3.5P,0.4V & 2.5L, 4.0P, 0.4V; LM visual cortex: 4.0L, 3.5P, 0.4V. Pulled glass capillaries were connected to a microinjection system (Stoelting Quintessential Stereotaxic Injector) to deliver viral vectors into the brain at a rate of 40–70 nL per minute, with a total volume of 100–200 nL per injection site. To express the calcium indicator GCaMP6s in neuronal cell bodies or long-range projection axons either AAV5-Syn-GCaMP6s or AAV1-Syn-GCaMP6s (1−10^13^ gc/mL; Addgene #100843) was injected into the relevant brain region. For experiments requiring expression of the soma-targeted GCaMP variant (riboGCaMP) in PM cell bodies, AAV9-Syn-GCaMP6m-RPL10a (1−10^13^ gc/mL; Addgene #158777) was injected into PM. For experiments requiring Cre-dependent optogenetic silencing of axon terminals within PM while imaging PM cell bodies, CAV2-Cre (1−10^13^ gc/mL; CRNS Plateforme de Vectorologie de Montpellier) was mixed in a 1:10 ratio with AAV5-syn-GCaMP6s and injected into PM and AAV5-Syn-SIO-eOPN3-mScarlet (2−10^13^ gc/mL; Addgene #125713) was injected into either LP, V1, or LM. All viruses were allowed to express for at least 3 weeks prior to experiments.

For headpost implantation, mice were anesthetized with isoflurane and the scalp was cleaned with Betadine solution. An incision was made at the midline and the scalp resected to each side to leave an open area of the skull. After cleaning the skull and scoring it lightly with a surgical blade, a custom titanium head post was secured with C&B-Metabond (Butler Schein) with the left PM or V1 centered. Two skull screws (McMaster-Carr) were placed at the right anterior and posterior poles (bilateral to the injection site). A 3x3-mm craniotomy was made over the left PM or V1. A glass window made of a 3x3-mm square inner cover slip adhered with an ultraviolet-curing adhesive (Norland Products) to a 5 mm-diameter round outer cover slip (both #1, Warner Instruments) was inserted into the craniotomy and secured to the skull with Cyanoacrylate glue (Loctite). A circular ring was attached to the titanium headpost with glue, and additional Metabond was applied to cover any exposed skull and to cover each skull screw. Analgesics were given immediately after surgery (5 mg/kg Carprofen and 0.05 mg/kg Buprenorphine) and on the two following days to aid recovery. Mice were given a course of antibiotics (Sulfatrim, Butler Schein) to prevent infection and were allowed to recover for 3-5 days following implant surgery before beginning wheel training.

### Histology

For histological preparation of brain sections, mice were deeply anesthetized with isoflurane and transcardially perfused with PBS followed by 4% PFA in PBS. Perfused brains were extracted and postfixed overnight at 4°C. 40-μm coronal sections were then prepared from the fixed brains using a vibratome (Leica VT1000 S). Free-floating sections were then mounted to microscope slides, coated with mounting medium (ProLong Gold Antifade Mountant with DAPI, Invitrogen) and sealed with cover glass. Epifluorescence and confocal images of brain sections were taken with a Zeiss Axio Imager 2 and Zeiss LSM 900, respectively.

### Calcium imaging

Calcium indicators in neuronal cell bodies and axon terminals within the cortex were imaged with a 2-photon Movable Objective Microscope (MOM) equipped with a galvo-resonant scanner (Sutter Instruments) through a 25x, 1.05 NA objective (Olympus). 2-photon excitation of the calcium indicators was achieved by scanning the output of a Ti-sapphire laser (MaiTai eHP DeepSee, SpectraPhysics) tuned to 920 nm. Emitted fluorescence was collected by a PMT (H10770PA-40, Hamamatsu) and digitized into image series with ScanImage software (Vidrio) at ∼30 Hz at a resolution of 256x256. To prevent light contamination from the display monitor, the microscope was enclosed in blackout material that extended to the headpost.

All calcium imaging was performed through a cranial window in awake, head-fixed mice that could freely run on a cylindrical wheel. Wheel speed was monitored by a magnetic angle sensor (Digikey). Before calcium imaging data were collected, mice were habituated to head-fixation for several days to ensure they spontaneously engaged in sustained locomotion bouts with normal posture. During calcium imaging experiments, the mouse’s face was illuminated with an IR LED array and facial video was acquired with a miniature CMOS camera (Blackfly s-USB3, Flir) at a frame rate of 10 Hz. All data acquisition signals (laser scanner frame times, wheel signal, and video camera frame times) were digitized at 5 kHz using a Power1401 data acquisition system (CED).

Imaging of cell bodies and axon terminals in layer 2/3 was performed at 150-350 μm depth relative to the brain surface, while imaging of apical dendrites in layer 1 was performed at 50-100 μm depth. For each mouse, 1-4 fields of view were imaged. During each session, spontaneous activity was collected for 10 mins before the series of visual stimuli were presented, and 10 mins after (20 mins total) as the mouse freely moved on the wheel in front of a mean-luminance gray screen.

### Optogenetic stimulation

For imaging sessions in which projection axons expressing the inhibitory opsin eOPN3 were optically stimulated, a two-session experimental design was used to compare conditions with and without optical stimulation. Since eOPN3 has a relatively long and potentially variable time constant for inactivation (∼5 min; Mahn et al., 2021), it is not well suited to traditional experimental designs using optogenetics in which control periods and periods with optical stimulation are interleaved throughout a session. Imaging data were therefore first collected during a session in which no optical stimuli were given (“Dark” session). After the end of this control session, the mouse was returned to its home cage for ∼2 hr. This waiting period ensured that residual effects of neuronal adaptation to visual stimulation did not carry over from one session to the next. After the waiting period, the mouse was placed back on the imaging set-up and the same field of view was imaged for a second session (“LED” session). During this session, light from a green LED (M530L3-530 nm, Thorlabs) was triggered with a 1-s TTL pulse, directed through the epifluorescence path of the Sutter MOM and reflected through the objective to produce widefield illumination onto the cranial window (∼1 mm spot diameter, ∼30 mW/mm^2^). This 1-s optical stimulus was given every 2 minutes throughout the session during intermittent pauses in scanning. Comparisons between neuronal properties were then made between the first and second imaging session to assess the effects of optogenetic manipulation.

### Visual stimulus presentation

Visual stimuli were presented on a 43 cm x 24 cm gamma-corrected LCD monitor (model B206HQL Aymph, Acer) positioned 45° from the mouse’s midline, with the center of the monitor located 18 cm from the mouse’s eye. The monitor subtended ∼100° azimuth/67° elevation of the mouse’s visual hemifield. A photodiode (model #SM1PD1B, Thorlabs) was fixed to the lower right corner of the monitor and connected to the Power1401 to allow precise recordings of visual stimulus onset times. Visual stimuli were generated and synchronized with the 60 Hz refresh rate of the monitor using MATLAB-based Psychtoolbox software (Brainard, 1997). Visual stimuli were 2 s in duration and consisted of either drifting sine-wave gratings of varying orientation, drift direction, size, spatial frequency, or temporal frequency or random dot kinematograms (RDKs). RDKs consisted of randomly positioned white circles (4° diameter) on an isoluminant gray background, such that 20% of the background was occupied by the circles. Each circle was then allowed to drift (80 °/s) in a random direction based on the value of percent motion coherence chosen for a given 2-s visual stimulus. For all types of visual stimuli, the time between the end of one visual stimulus and the onset of the next (i.e. inter-stimulus interval) was 5 s.

## DATA ANALYSIS

Data were analyzed using custom-written scripts in Mathematica 11 (Wolfram Research) and MATLAB R2018a (Mathworks).

### Cell counts and fluorescence intensity quantification in histological sections

Fluorescently labeled cells were counted manually in each histological section. Sections were aligned to bregma based on the detection of a small volume (<50 nL) of red retrobeads (Lumafluor) injected into the brain at the anterior-posterior coordinate of bregma immediately before transcardial perfusions as a fiducial marker. Fluorescently labeled cells were assigned to specific brain regions based on alignment with coronal sections in the Paxinos-Franklin mouse brain atlas (Paxinos and Franklin, 2008). To calculate the total cell density in each brain region, the total number of cells counted in that region was divided by an estimate of its total volume, computed by numerical integration of the cross-sectional areas of the portions of each section corresponding to the brain region. Normalized cell count as a function of anterior-posterior distance from bregma was calculated by dividing the cell count associated with a given histological section by the highest cell count among all the sections from a given animal. Fluorescence intensity of axonal or dendritic labeling was measured in 10- or 5-μm bins across the laminar depth of the cortex relative to the pia for whole-cortex and layer 1 measurements, respectively. Mean normalized laminar fluorescence intensity for a given animal was calculated by dividing all fluorescence intensities by the highest fluorescence intensity for that animal and then averaging resulting normalized fluorescence vs. depth data from all sections in that animal.

### Wheel position and change-points

Wheel position was determined from the output of the linear angle detector. The circular wheel position variable was first transformed to the [-ρε, ρε] interval. The phases were then circularly unwrapped to yield running distance as a linear variable, and locomotion speed was computed as a differential of distance (cm/s). A change-point detection algorithm detected locomotion onset/offset times based on changes in standard deviation of speed. Locomotion onset or offset times were estimated from periods when the moving standard deviations, as determined in a 0.5-s window, exceeded or fell below an empirical threshold of 0.1. Locomotion trials were required to have average speed exceeding 0.5 cm/s and last longer than 1 s. Quiescence trials were required to last longer than 2 s and have an average speed < 0.5 cm/s.

### Quantification of calcium signals

Analysis of imaging data was performed using ImageJ and custom routines in MATLAB (Mathworks). Motion artifacts and drifts in the Ca^2+^ signal were corrected with the moco plug-in in ImageJ (Dubbs et al., 2016), and regions of interest (ROIs) were selected as previously described (Chen et al., 2013). All pixels in each ROI were averaged as a measure of fluorescence, and the neuropil signal was subtracted (Chen et al, 2013; Lur et al., 2016; Tang and Higley, 2020; Ferguson et al., 2023). ΔF/F was calculated as (F-F_0_)/F_0_, where F_0_ was the lowest 10% of values from the neuropil-subtracted trace for each session.

For some analyses (Figure 4 and Figure S6), ΔF/F signals were deconvolved to estimate the times of underlying Ca^2+^ events. The AR1 FOOPSI algorithm in the Oasis toolbox (Friedrich et al., 2017) was used to find the optimal convolution kernel, baseline fluorescence, and noise distribution. Ca^2+^ events were then detected as times at which the deconvolved signal increased beyond 3 standard deviations. Time-varying Ca^2+^ event rates were then computed as

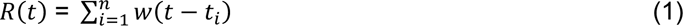

where *t_i_* indicates the time at which a Ca^2+^ event occurred for *n* events in a recording, and *w* is a sliding Gaussian window,

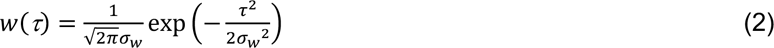

where α*_w_* = 100 ms.

### Cellular ROI selection

Image series collected from 2-photon calcium imaging were corrected for motion artifacts using the moco plug-in on ImageJ (Dubbs et al., 2016). ROIs corresponding to cell bodies or axon terminals within the imaging field of view were chosen with a semi-automated procedure using mean- or max-projections of the imaging series (Chen et al., 2013). All pixels within an ROI were averaged as a measure of fluorescence. To correct for fluorescence originating from neuropil potentially contaminating the ROI signal, a group of pixels corresponding to a ∼20 μm region around each ROI (excluding all selected ROIs) was averaged and subtracted from the ROI signal at each time point:

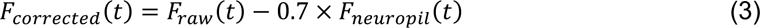

From these neuropil-corrected raw fluorescence traces for each ROI, calcium activity was quantified as ττ*F/F*:

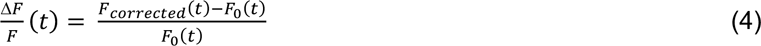

where *F*_0_(*t*) is a sliding 10^th^ percentile of the raw fluorescence distribution, with a sliding window 60 s in width.

For ROIs corresponding to axon terminals, additional efforts were made to minimize inclusion of redundant ROIs that putatively belong to the same neuron in the presynaptic source region. Whereas redundancies in cell body ROIs are highly unlikely (i.e. 2 or more ROIs corresponding to the same neuronal cell body), ROIs for axon terminals could suffer from redundancy due to axonal branching. In order to minimize the number of redundant axon terminal ROIs in our dataset, we used a procedure based on activity correlations. For this procedure, we first generated a dataset derived from axonal calcium imaging sessions consisting of axon segments that were unambiguously part of the same stretch of axon (“sub-ROIs”). For all such sub-ROI pairs within an imaging field of view, we then calculated the partial correlation of their activity throughout the entire imaging session, subtracting the mean value of all pairwise correlation values from each raw pairwise correlation. The resulting distribution of partial correlation values corresponds to the case in which all axon terminal ROIs in a dataset are actually from the same neuron. We then compared this distribution of sub-ROI partial correlation values with that of the partial correlation values between axon terminal ROIs that were considered to belong to different neurons based upon structural images (i.e. terminals that were non-contiguous within the field of view). Based on an analysis of the distributions of correlations for ROIs and sub-ROIs (Figure S1), a partial correlation value of 0.3 was the most conservative threshold for separating these distributions in our datasets. Thus, in our larger dataset of axon terminal ROIs, if any 2 ROIs exhibited partial correlations ≥ 0.3, one of the ROIs was discarded as putatively redundant and not included in further analysis. This method was also applied to eliminate putatively redundant ROIs from experiments in which we imaged apical dendrites from pyramidal neurons arborizing in layer 1 (Figure S1, Figure S4).

### Quantification of visual responses and visual feature selectivity

Prior to imaging sessions used for characterizing neuronal visual responses, the population receptive field for the ROIs in a field of view was mapped by partitioning the LCD monitor into a 3x3 grid and presenting binarized Gaussian noise stimuli (Niell and Stryker, 2008) within each patch in the grid. In subsequent imaging experiments using a given field of view, the monitor was positioned such that it was centered at the location within the visual field that elicited the largest visual responses from the neuron ROIs during these initial receptive field mapping sessions. With the exception of size tuning and motion coherence protocols (see below), visual stimuli were ∼40° in diameter.

For ROIs corresponding to cell bodies or axon terminals, visually evoked response magnitude was quantified as the mean z-scored Δ*F/F* during a 2-s visual stimulus, with the mean and standard deviation of the Δ*F/F* signal during a 1-s baseline period prior to visual stimulus onset used for z-scoring. The overall visual response magnitude for a given ROI was the mean visually evoked z-scored Δ*F/F* averaged across all visual stimulus presentations during an imaging session. ROIs were considered significantly visually responsive if Δ*F/F* values during visual stimulation periods were significantly larger than Δ*F/F* values during 1-s periods before visual stimulus onset (*p* < 0.05, Student’s t-test). Calculation of selectivity for different visual features is described below.

### Orientation/direction selectivity

To calculate an ROI’s selectivity for the orientation and direction of drifting sine-wave gratings, 12 different drift directions were presented during imaging sessions (evenly spaced between 0° and 352°). The temporal frequency of the gratings was 1 Hz and the spatial frequency was either 0.05, 0.085, or 0.12 cycles/°. Each combination of drift direction, temporal frequency, and spatial frequency was randomly repeated 10x during an imaging session. For each ROI, orientation selectivity was quantified by *L_ori_* = [1 – Circular Variance] (Mazurek et al., 2014):

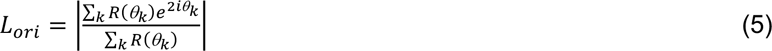

where *R*(8*_k_*) is the mean visual response magnitude to drift direction 8*_k_*, and direction selectivity was quantified by *L_dir_* = [1 – Directional Circular Variance]:

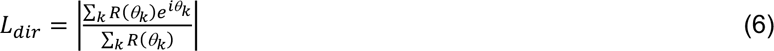

### Spatial frequency, temporal frequency, & speed tuning

To calculate an ROI’s tuning for visual spatial frequency (SF), temporal frequency (TF), and speed, the spatial and temporal frequencies of sine-wave gratings drifting at 0° were varied approximately in octaves (SF: 0.02, 0.04, 0.08, 0.16, 0.32 cycles/°; TF: 0.5, 1, 2, 4, 8, 15, 24 Hz) and each SF-TF combination was randomly repeated 15x during an imaging session. Each ROI’s response to a specific SF or TF was computed as the mean visual response magnitude for each presentation of that SF or TF. Different SF-TF combinations corresponded to different speeds (in °/s), which was calculated as the ratio between TF and SF. Each ROI’s response to a specific speed was thus computed as the mean visual response magnitude for a unique ratio between TF and SF. To quantify the preferred SF, TF, and speed for a given ROI, visual responses as a function of either SF, TF, or speed were fit to log-Gaussian models (Priebe et al., 2006):

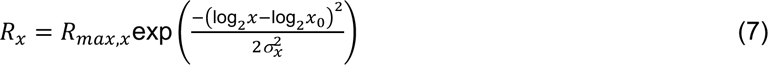

where *x* is either SF, TF, or speed and *R_max,x_*, *x*_0,_ and α*_x_* are parameters fit by nonlinear optimization. SF, TF, and speed tuning curves were considered well-fit if the model *R^2^*was greater than 0.8. In this model, *x*_0_ corresponds to preferred SF, TF, or speed.

### Size tuning

To calculate an ROI’s tuning for visual stimulus size, the diameter of drifting sine-wave gratings within the population receptive field was varied between 7° and 75° in increments of 10°. The temporal frequency of the gratings was 1 Hz, the spatial frequency was either 0.05, 0.085, or 0.12 cycles/°, and the drift direction was in either of the 4 cardinal directions. Each combination of size, temporal frequency, spatial frequency, and drift direction was randomly repeated 4x during an imaging session. The amount of surround suppression exhibited by an ROI was calculated from its size-tuning curve as a suppression index (*SI*):

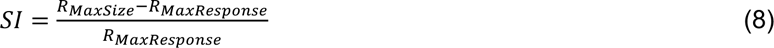

where *R_MaxSize_* is the magnitude of the response for the maximum size in stimulus set (75°) and *R_MaxResponse_* is the magnitude of the maximum response across all stimulus sizes.

### Motion coherence selectivity

To characterize an ROI’s selectivity for coherent motion within the visual field, full-screen random dot kinematograms (RDKs) (see “Visual stimulus presentation”) were presented in which the percent motion coherence was varied among 8 values (20, 25, 31, 40, 50, 63, 79, 100%). For each percent motion coherence value, drift direction could be in either of the 4 cardinal directions. Each combination of percent motion coherence and drift direction was randomly repeated 15x during an imaging session. For each ROI, the drift direction that elicited the largest visual responses was chosen as that ROI’s preferred direction and visual responses as a function of motion coherence were analyzed for that drift direction. To quantify an ROI’s selectivity for coherent visual motion, the Pearson correlation coefficient between a vector of visual responses for each motion coherence value and a vector of the corresponding motion coherence values was calculated (Sit and Goard, 2020).

### Quantification of modulation of neuronal activity by locomotion

For calculation of locomotion modulation index (LMI), only locomotion bouts occurring after and before at least 15 s of quiescence were used. LMI for each ROI was calculated as

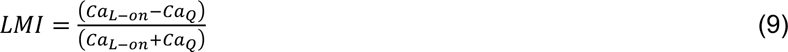

where *Ca_L-on_* is the mean Δ*F/F* during the 4 s following locomotion onset and *Ca_Q_* is the mean Δ*F/F* during the 10–15 s after locomotion offset.

### Quantification of modulation of neuronal activity by pupil size

From video of the mouse’s face, pupil size of the non-visually-stimulated eye was computed from individual video frames. Frames were first cropped to a region around the eye and image pixels were binarized such that the majority of the pixels within the pupil were white and the majority of the pixels outside of the pupil were black (the IR light from the Ti-sapphire laser that escapes through the mouses eye causes the pupil to appear white when captured by the camera). Canny edge detection was then used on the binarized image frames to detect edges of the pupil. Pupil size was then estimated as the authalic diameter of the area enclosed by the detected edges (i.e. the diameter of a circle with the same area) (Neske et al., 2019).

All analyses relating neuronal activity to pupil fluctuations were restricted to quiescent (i.e. non-locomotion) periods. To relate pupil dilation and constriction to fluctuations in neuronal activity, the time series of pupil size was bandpass-filtered between 0.1 and 1 Hz. The bandpass-filtered signal of pupil size was converted to phase of the pupil dilation-constriction cycle through a Hilbert transform, which transforms the signal into a time series of complex numbers. The inverse tangent of the ratio between the imaginary and real parts of the complex number at each time point yields the phase of the bandpass-filtered pupil signal. To quantify neuronal activity as a function of the phase of the dilation-constriction cycle, Δ*F/F* time series for each ROI were also bandpass-filtered between 0.1 and 1 Hz. These bandpass-filtered Δ*F/F* signals were then z-scored using baseline periods associated with minimal changes in pupil size and small baseline pupil diameter. Specifically, these baseline periods needed to be associated with a mean of the bandpass-filtered pupil signal that was less than 1.5x the interquartile range of this signal *and* a mean pupil size that was within the bottom 25^th^ percentile of the distribution of pupil sizes observed during quiescent periods during a given session. The mean and standard deviation of the bandpass-filtered Δ*F/F* signal during these baseline periods was used for z-scoring the Δ*F/F* signal. For each dilation-constriction cycle, z-scored Δ*F/F* for each ROI was then averaged in bins of width π/32 from −π to π. Only dilation-constriction periods larger than a threshold of 1.5x the interquartile range of the bandpass-filtered pupil signal were analyzed.

To calculate cross-correlations between neuronal activity and pupil fluctuations, both the time series of pupil size and the time series of Δ*F/F* for each ROI were lowpass-filtered at 10 Hz. Correlations between these signals were then computed at different lags in 100-ms increments in a time window of 8 s, using the pupil signal as the reference signal. The strength of the modulation of Δ*F/F* by pupil size for each ROI was quantified as the value of the peak cross-correlation between Δ*F/F* and pupil size.

### Quantification of modulation of neuronal activity by facial motion/whisking

From video of the mouse’s face, a region consisting of the nose, mouth, and whisker pads was cropped from each frame. A region of interest was then drawn around the whisker pad and facial motion energy was calculated as the absolute value of the difference in average pixel intensity of the whisker pad between successive image frames. Onset times of facial motion bouts were computed by first z-scoring the entire time series of facial motion energy. High and low thresholds of facial motion energy were then defined as the 60% and 40% quantiles of the z-scored facial motion energy distribution during quiescent periods. After smoothing the time series of z-scored facial motion energy with a 1-s moving-average filter, high and low facial movement states were defined as periods in which the smoothed signal remained above or below the high and low threshold levels for at least 500 ms. Onset of a facial motion bout was then defined as the time at which a high facial motion state was preceded by at least 4 s of a low facial motion state. ROI calcium activity aligned to these onsets was z-scored using the mean and standard deviation of the Δ*F/F* during low facial motion periods.

To calculate cross-correlations between neuronal activity and facial motion, both the time series of facial motion energy and the time series of Δ*F/F* for each ROI were lowpass-filtered at 10 Hz. Correlations between these signals were then computed at different lags in 100-ms increments in a time window of 8 s, using the facial motion energy as the reference signal. The strength of the modulation of Δ*F/F* by facial motion for each ROI was quantified as the value of the peak cross-correlation between Δ*F/F* and facial motion energy.

### Calculation of Ca^2+^ event-triggered averages

For data shown in Figure 4E,F and Figure S6B,C, Ca^2+^ activity aligned to cell body Ca^2+^ events was computed as follows. For each riboGCaMP6m-expressing cell body identified in a field of view, the times of its Ca^2+^ events were estimated from the deconvolved ΔF/F signal. In an 8-s window around each Ca^2+^ event from the cell body, the population average Ca^2+^ event rate from all other relevant ROIs (i.e. axon ROIs or ROIs from other cell bodies) was calculated. For each cell body whose event-triggered average was calculated this way, the peak event rate was taken as the maximum event rate in the 8-s window.

### Statistical tests

In both the text and figures “*N*” refers to number of mice and “*n*” refers to number of neuron ROIs (cell bodies or axon terminals). Unless otherwise noted, data are summarized by either box-and-whisker charts (with “boxes” marking the median value and the 25% and 75% quantiles, and “whiskers” marking the minimum and maximum values in the dataset) or summary curves indicating the mean value and shading indicating the standard error of the mean.

For statistical comparisons among groups, we accounted for the nested structure of the data (i.e. cellular ROIs nested within mice). Without taking nesting into consideration, false positive rates are inflated since cellular ROIs from the same mouse cannot be considered independent measurements (Aarts et al., 2014). We used semi-weighted error estimators to account for the nesting of cellular ROIs within mice (Lur et al., 2016). For this procedure, let *y_i_* be the mean for a given parameter (e.g. locomotion modulation index) for the *i*-th mouse in a group, where *x_j_* is the parameter value for the *j*-th ROI in a set if *M* ROIs for mouse *i*:

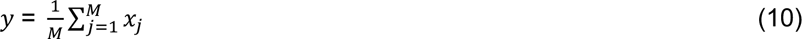

The unweighted estimator of the mean is defined as the mean of *y_i_* for all mice in a group of *N* mice, where *N* is the number of degrees of freedom. This analysis is suboptimal, as some animals have many cells and yield more reliable estimates, whereas other animals have few cells and yield less reliable estimates. At the other extreme, the weighted estimator is defined by pooling parameter values from all ROIs across the *N* mice in a group and using this number as the degrees of freedom. While the weighted estimator is commonly used in statistical comparisons, the false positive rate is improperly controlled in this case. The semi-weighted estimator for a given parameter measured from multiple ROIs from multiple mice in a group is defined as:

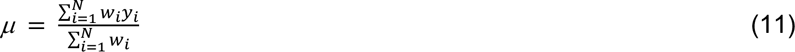

where the weight for the *i*-th mouse is defined as

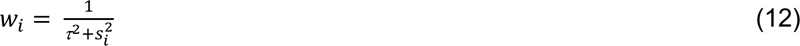

and *s_i_^2^* is the within-mouse variance across ROIs and τ^2^ is the estimated variance across all mice within the group using maximum likelihood estimation (Chung et al., 2013):

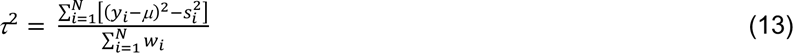

Equations 12 and 13 were solved iteratively and checked for convergence of τ^2^ to <0.001. If there is minimal across-mouse variance, then the contribution of each ROI is weighted by the number of mice (the weighted estimator), whereas if across-mouse variance is large (i.e. parameter measurements for ROIs within a mouse are strongly dependent), then each mouse is given the same weight.

The semi-weighted estimators derived for different cell types (e.g. V1 axons vs. LP axons) or experimental conditions (e.g. control vs. eOPN3) were then used to calculate the statistical significance of pairwise comparisons using a Student’s t-test. The Benjamini-Hochberg procedure was used to control the false discovery rate from multiple pairwise comparisons. *P*-values < 0.05 after multiple comparisons correction were considered statistically significant. Central markers on box-whisker plots indicate median values, boxes delineate the 25^th^ and 75^th^ percentiles, and upper/lower whiskers indicate maximum/minimum values.

## Acknowledgements

The authors thank all members of the Higley and Cardin laboratories for helpful input throughout all stages of this study. We thank Rima Pant for generation of AAV vectors. We thank Dr. Ofer Yizhar for the gift of the eOPN3 plasmid. This work was supported by funding from the NIH (R01EY022951 R01MH113852 to JAC, F32NS100279 and K99EY030550 to GTN, EY026878 to the Yale Vision Core), an award from the Kavli Institute of Neuroscience (to JAC), and support from the Ludwig Foundation (to JAC).

## Author Contributions

GTN and JAC designed the experiments. GTN collected and analyzed the data. GTN and JAC wrote the manuscript.

---

**Supplementary Figure 1.**
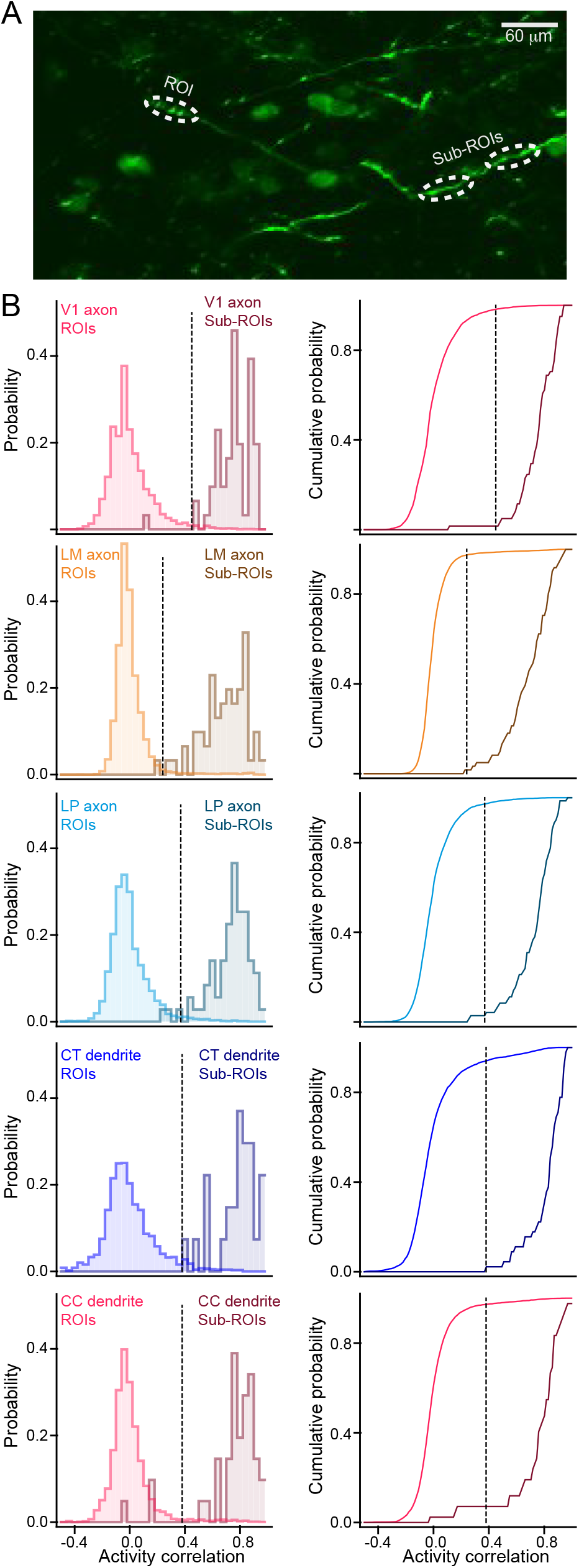
Correlation-based curation of axon and dendrite ROIs. A. Example of defining “ROIs” and “sub-ROIs,” the latter of which are 2 fluorescent segments that unambiguously belong to the same axon or dendrite. The distribution of activity correlation values between pairs of sub-ROIs serves as a null distribution for deciding whether any 2 chosen axon/dendrite ROIs originate from different neurons (see also Materials and Methods). B. Comparisons of activity correlations between ROIs and subROIs for all axon and dendrite types studied. Left: Probability histograms of activity correlations for ROIs and sub-ROIs. Dashed lines indicate criteria for determining redundancy in axon/dendrite ROIs: any 2 axon/dendrite ROIs in the dataset exhibiting an activity correlation above the criterion are considered redundant, and only one is kept for further analysis. Right: Cumulative probability histograms of activity correlations for ROIs and sub-ROIs. Dashed lines correspond to the maximum difference between ROI and sub-ROI cumulative probability histograms and serve as the criteria for determining axon/dendrite redundancy. (V1→PM axons: N = 10 animals, nROIs = 3437, nSub-ROIs = 61; LM→PM axons: N = 8 animals, nROIs = 6927, nSub-ROIs = 61; LP→PM axons: N = 9 animals, nROIs = 3857, nSub-ROIs = 71; V1 CT dendrites: N = 4 animals, nROIs = 3864, nSub-ROIs = 45; V1 CC dendrites: N = 4 animals, nROIs = 4207, nSub-ROIs = 42).

**Supplementary Figure 2.**
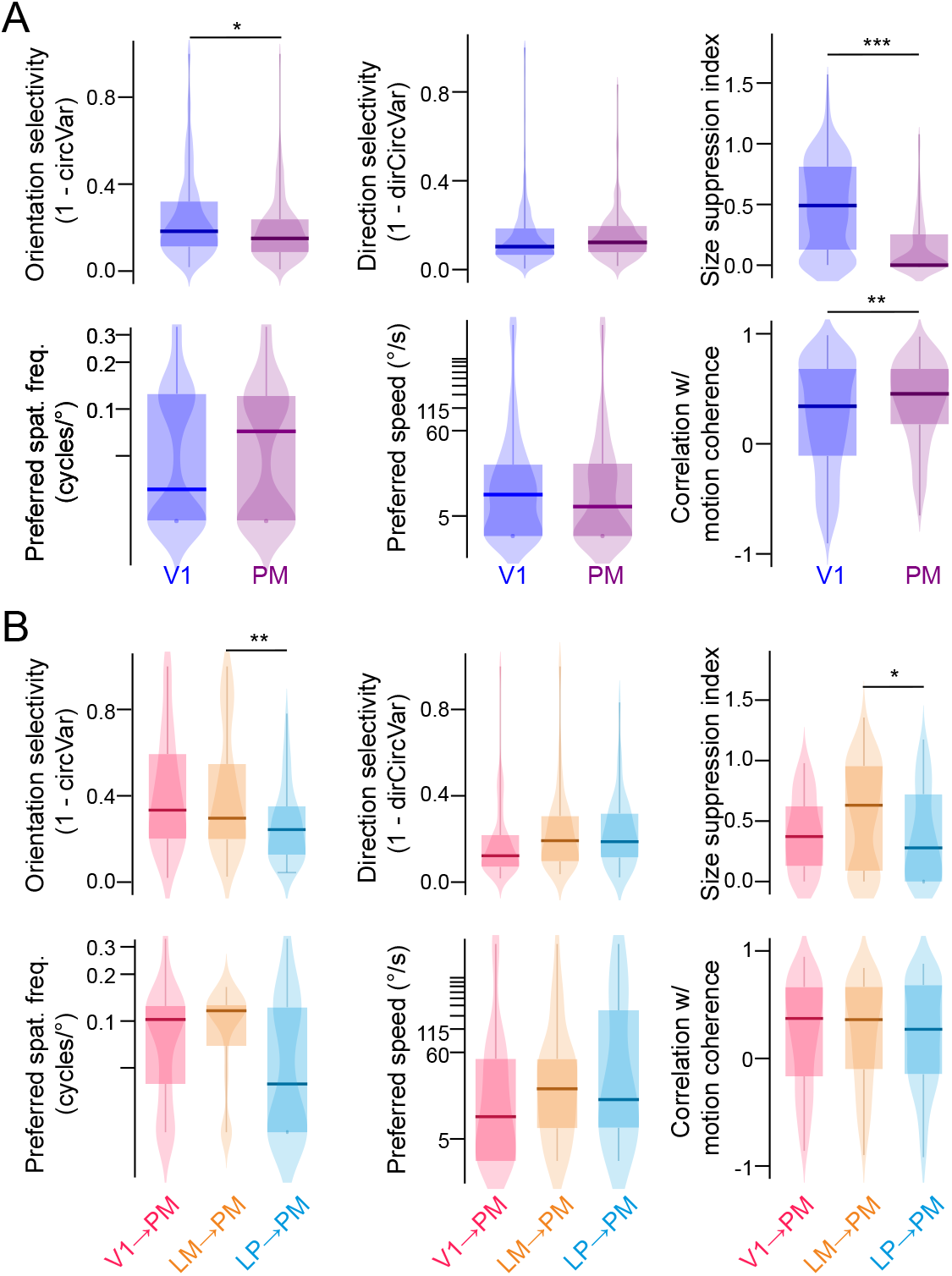
Visual feature selectivity of PM and V1 neurons and corticocortical and higher-order thalamocortical projections to PM. A. Quantification of orientation/direction selectivity, surround suppression, preferred visual spatial frequency/speed, and sensitivity to motion coherence in V1 and PM cell bodies. B. As in A, but for V1→PM, LM→PM, and LP→PM axons. Sample sizes and p-values are found inSupplementary Table 1. *p<0.05, **p<0.01, ***p<0.001, semi-weighted t-test, Benjamini-Hochberg correction for false discovery rate.

**Supplementary Figure 3.**
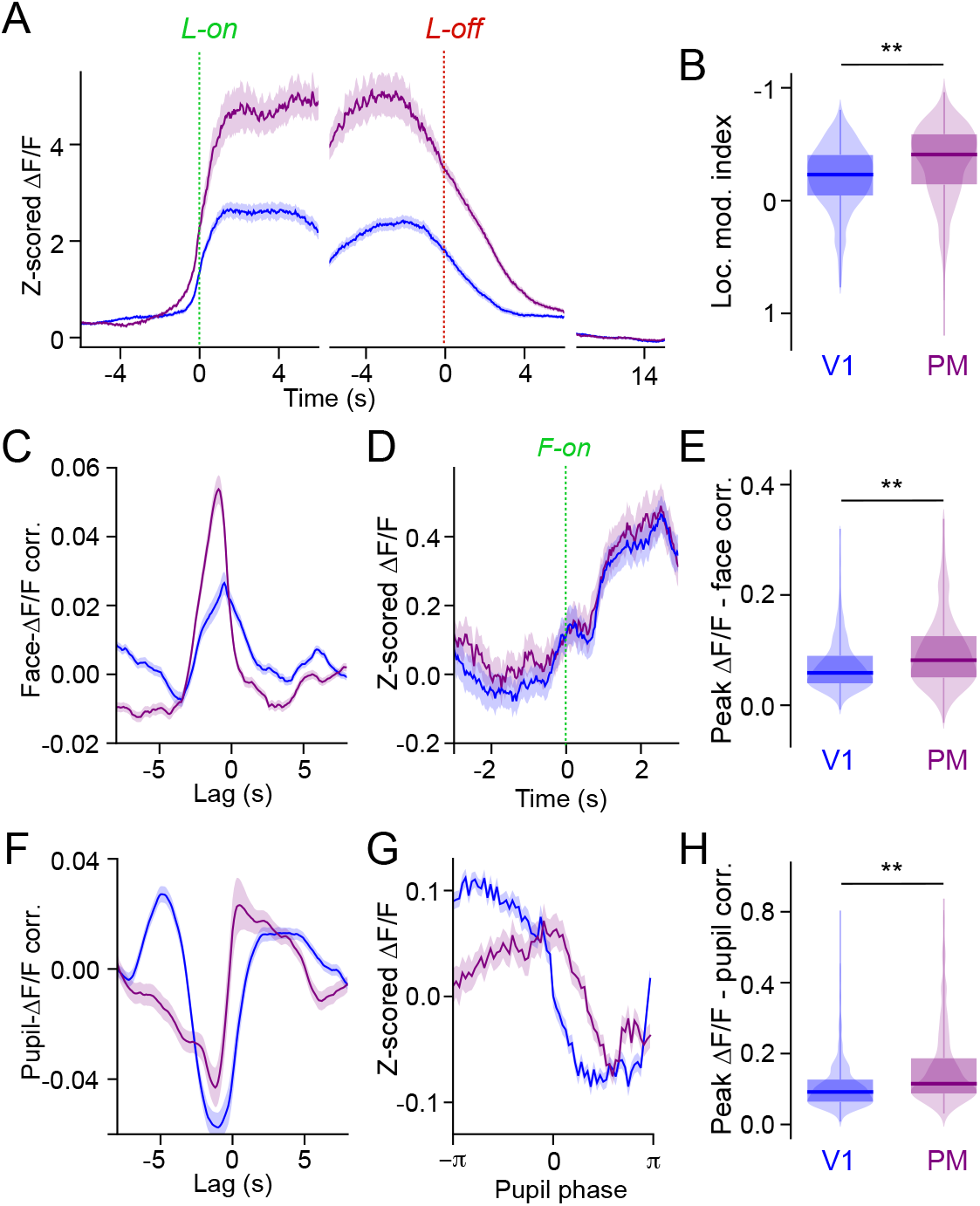
State modulation of V1 and PM neurons. A. Neuronal activity (Z-scored ΔF/F) aligned to locomotion onset and offset. Baseline activity (10-15 s after locomotion offset during sustained quiescent periods) is shown for comparison. B. Locomotion modulation indices for ROIs of each cell type. C. Cross-correlation between neuronal activity (ΔF/F) and facial motion (whisker pad motion energy). Facial motion is the reference signal in the cross-correlation (i.e. a peak at negative lag values indicates that neuronal activity increases after a corresponding increase in facial motion). D. Neuronal activity (Z-scored ΔF/F) aligned to the onset of high facial motion periods. E. Peak correlation values between neuronal activity and facial motion for ROIs of each cell type. F. Cross-correlation between neuronal activity (ΔF/F) and pupil size. G. Neuronal activity (Z-scored ΔF/F) aligned to one pupil dilation-constriction cycle (derived from the Hilbert transform of the pupil signal). H. Peak correlation values between neuronal activity and pupil size for ROIs of each cell type. Sample sizes and p-values are found in Supplementary Table 1. *p<0.05, **p<0.01, ***p<0.001, semi-weighted t-test, Benjamini-Hochberg correction for false discovery rate. Error bars denote s.e.m.

**Supplementary Figure 4.**
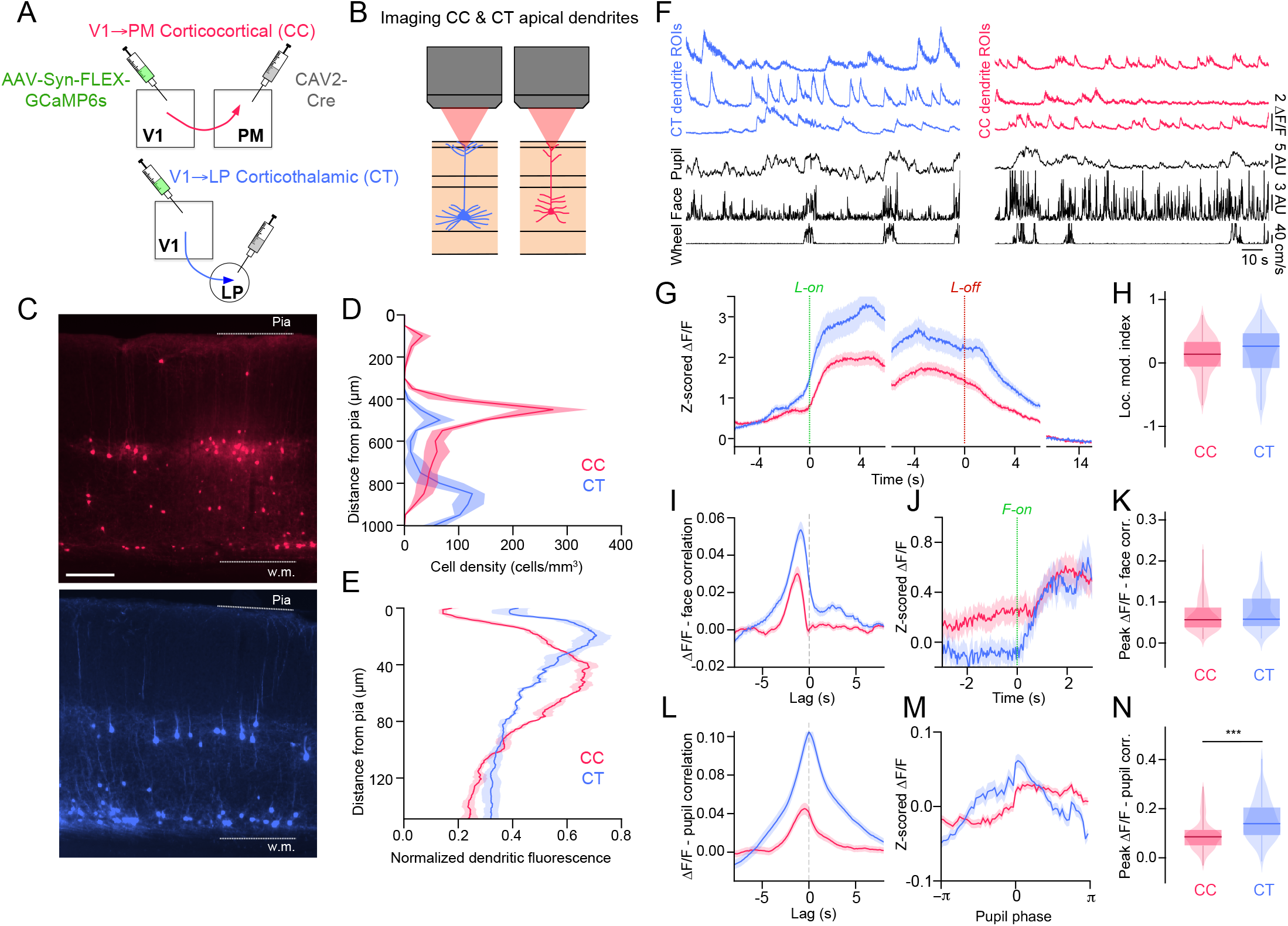
State modulation of layer 5 corticocortical and corticothalamic projection neurons in V1. A. Schematic of injections for labeling different V1 neurons based on projection target (PM and LP). B. Imaging apical dendrites of layer 5 projection neurons arborizing in layer 1. C. Fluorescence images of GCaMP6s expressed in V1→PM corticocortical projection neurons (top) and V1→LP corticothalamic projection neurons (bottom). D. Quantification of V1→PM corticocortical (CC) and V1→LP (CT) cell density along the laminar depth of V1. E. Laminar distribution of fluorescence from dendritic arbors in V1 layer 1. F. Fluorescence traces of dendritic Ca2+ activity from CC and CT projection neurons, simultaneous with behavioral state monitoring (locomotion speed, facial motion, pupil size) G. Neuronal activity (Z-scored ΔF/F) aligned to locomotion onset and offset. Baseline activity (10-15 s after locomotion offset during sustained quiescent periods) is shown for comparison. H. Locomotion modulation indices for ROIs of each cell type. I. Cross-correlation between neuronal activity (ΔF/F) and facial motion (whisker pad motion energy). Facial motion is the reference signal in the cross-correlation (i.e. a peak at negative lag values indicates that neuronal activity increases after a corresponding increase in facial motion). J. Neuronal activity (Z-scored ΔF/F) aligned to the onset of high facial motion periods. K. Peak correlation values between neuronal activity and facial motion for ROIs of each cell type. L. Cross-correlation between neuronal activity (ΔF/F) and pupil size. M. Neuronal activity (Z-scored ΔF/F) aligned to one pupil dilation-constriction cycle (derived from the Hilbert transform of the pupil signal). N. Peak correlation values between neuronal activity and pupil size for ROIs of each cell type. Sample sizes and p-values are found in Supplementary Table 1. *p<0.05, **p<0.01, ***p<0.001, semi-weighted t-test, Benjamini-Hochberg correction for false discovery rate. Error bars denote s.e.m.

**Supplementary Figure 5.**
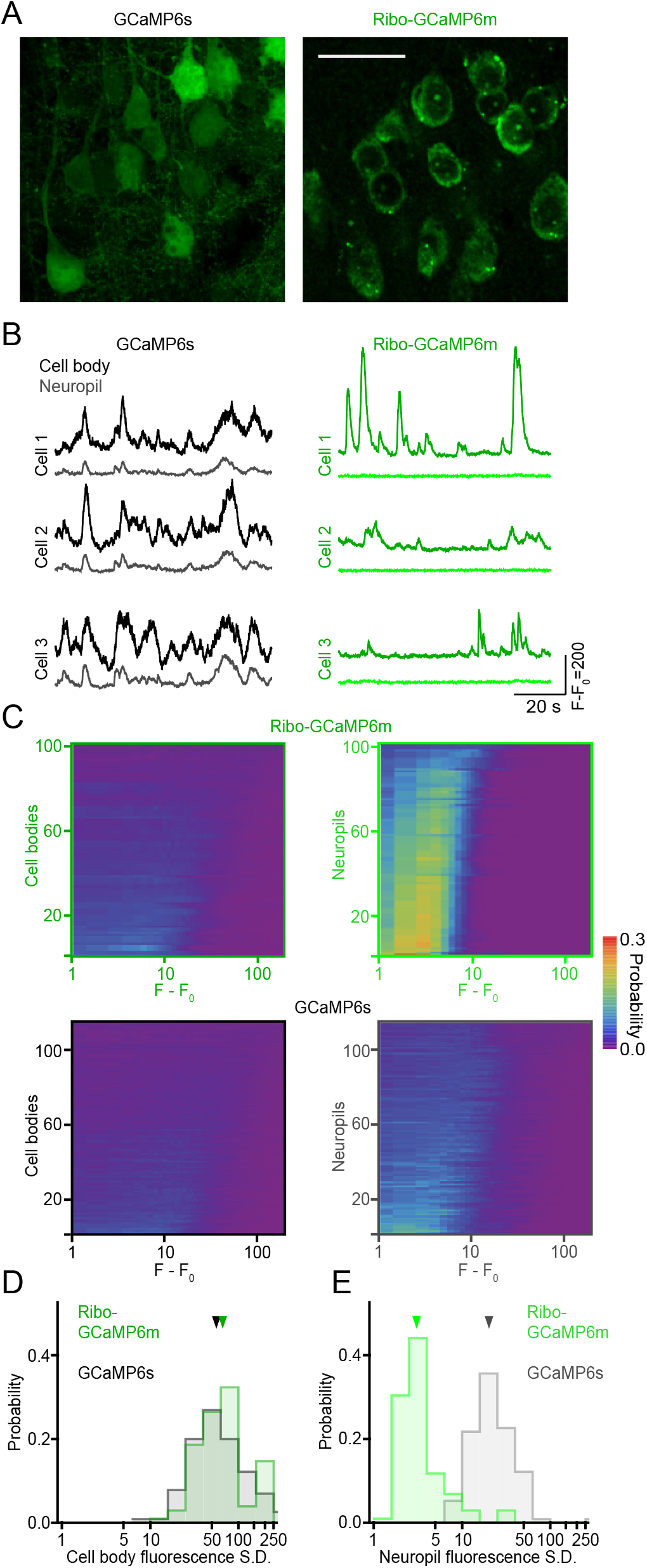
Negligible neuropil labeling using riboGCaMP6m. A. Comparison of GCaMP6s (left) and riboGCaMP6m (right) when expressed in PM cortical neurons. Scale bar = 30 μm. B. In vivo fluorescence signal (F-F0) from cell bodies and neuropil using GCaMP6s (left) and riboGCaMP6m expression (right) in PM cortical neurons. C. Density plots representing probability histograms of F-F0 values for cell bodies (right) and associated neuropil (left) for PM neurons expressing GCaMP6s and riboGCaMP6m. Each horizontal slice of a density plot is a probability histogram for an individual cell body/neuropil region. D. Probability histograms of the standard deviation (S.D.) of the fluorescence signal from PM neuron cell bodies expressing GCaMP6s or riboGCaMP6m. E. Probability histograms of the standard deviation (S.D.) of the fluorescence signal from the neuropil regions around PM neuron cell bodies expressing GCaMP6s or riboGCaMP6m. (GCaMP6s data: N = 3 animals, n = 115 ROIs; riboGCaMP6m data: N = 2 animals, n = 102 ROIs).

**Supplementary Figure 6.**
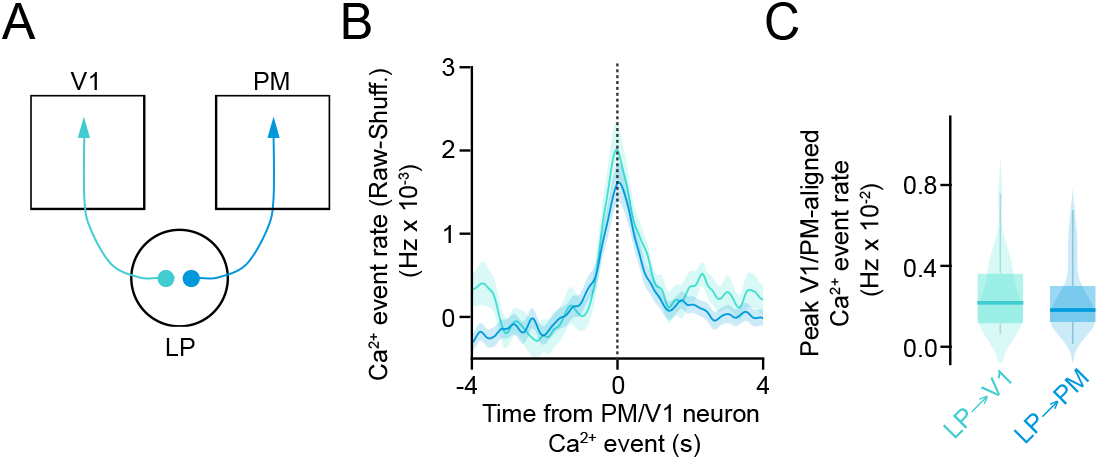
Comparison of LP projections to V1 and PM neurons. A. Schematic of two thalamocortical pathways from LP: one projecting to V1 (a canonical feedback, modulatory pathway) and one projecting to PM (the focus of the present study). B. Ca^2+^ event rates in LP axons aligned to Ca^2+^ events in PM/V1 neurons (raw data minus data from shuffled PM/V1 event times). For each PM/V1 neuron in a field-of-view, the mean Ca^2+^ event rate for all neighboring LP axon terminals was calculated in an 8-s window around the time of Ca^2+^ event onset in that PM/V1 neuron. C. Peak LP axon Ca^2+^ event rates aligned to PM/V1 neuron Ca^2+^ events. (LP→PM activity aligned with PM events: N = 5 animals, n = 56 PM neuron ROIs; LP→V1 activity aligned with V1 events: N = 3 animals, n = 23 V1 neuron ROIs). Error bars denote s.e.m.

**Supplementary Figure 7.**
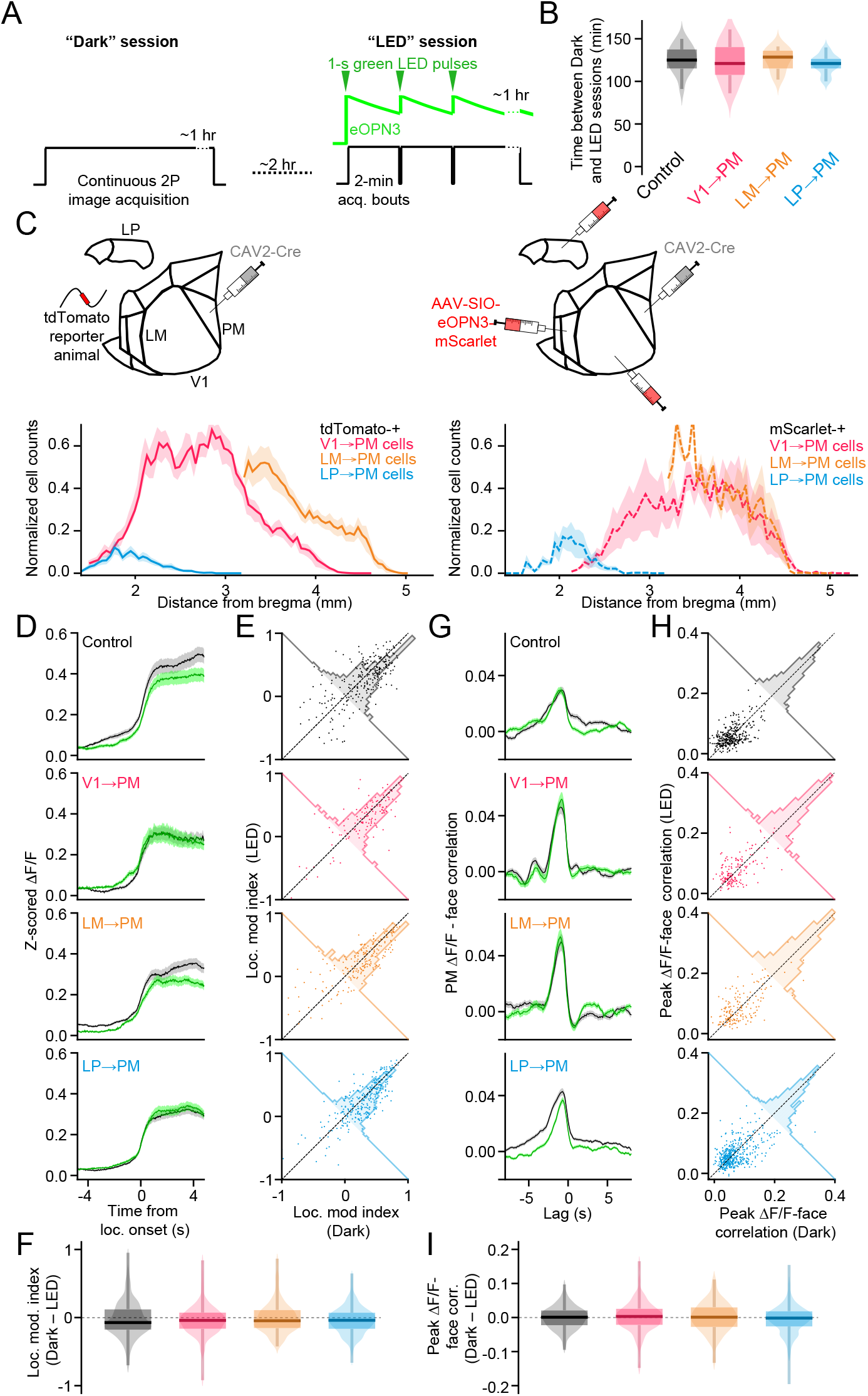
Further analyses of eOPN3 experiments. A. Schematic of the optogenetic stimulation strategy. During one imaging session (“Dark”), 2-photon imaging acquisition is continuous and no optogenetic stimuli are given. After a ∼2-hr waiting period, a second session (“LED”) is run for the same field of view, in which 2-min 2-photon imaging acquisition bouts are interrupted (∼5 s) by the application of 1-s green LED pulses reflected through the microscope objective onto the cranial window. The long time constant of eOPN3 activation (∼5 min) allows for continued suppression of synaptic release after the LED stimuli are applied. B. Verification that the time interval between “Dark” and “LED” imaging sessions did not systematically differ among experimental groups. (Control: N = 7 animals, n = 15 Dark-LED session pairs; V1→PM eOPN3: N = 7 animals, n = 17 Dark-LED session pairs; LM→PM eOPN3: N = 4 animals, n = 10 Dark-LED session pairs; LP→PM eOPN3: N = 6 animals, n = 21 Dark-LED session pairs). C. Quantification of the relative numbers of cells labeled with AAV injections of Cre-dependent eOPN3 after CAV2-Cre injection in PM. Retro-labeled cell counts from tdTomato reporter animals injected with CAV2-Cre in PM (Left, data from main Fig. 1) represent the “ground truth” of the relative numbers of V1, LM, and LP neurons that project to PM. (Right) Cell counts from individual (non-reporter) animals injected with CAV2-Cre in PM followed by injections of AAV-SIO-eOPN3-mScarlet in V1, LM, and LP in the same animals with injection volumes used for labeling these structures individually (main Fig. 5). (Ai9 reporter: N = 10 animals; eOPN3-injected: N = 3 animals). D. Neuronal activity (Z-scored ΔF/F) aligned to locomotion onset in control animals and animals with eOPN3 expressed in corticocortical and higher-order thalamocortical axons for session with (“LED”) and without (“Dark”) optogenetic stimulation. E. Locomotion modulation indices from individual PM neurons identified during both “Dark” and “LED” imaging sessions. F. Differences between locomotion modulation indices during sessions with (“LED”) and without (“Dark”) optogenetic stimulation. (Control: N = 7 animals, n = 252 ROIs; V1→PM eOPN3: N = 4 animals, n = 127 ROIs; LM→ PM eOPN3: N = 3 animals, n = 200 ROIs; LP→PM eOPN3: N = 5 animals, n = 292 ROIs). G. As in D but for cross-correlations between PM neuronal activity and facial motion. H. As in E but for peak correlation values between neuronal activity and facial motion. I. As in F but for differences in peak ΔF/F-face correlation values between “Dark” and “LED” sessions. (Control: N = 7 animals, n = 327 ROIs; V1→PM eOPN3: N = 4 animals, n = 169 ROIs; LM→PM eOPN3: N = 3 animals, n = 167 ROIs; LP→PM eOPN3: N = 5 animals, n = 468 ROIs). *p<0.05, **p<0.01, ***p<0.001, semi-weighted t-test, Benjamini-Hochberg correction for false discovery rate.

**Supplementary Table 1.**
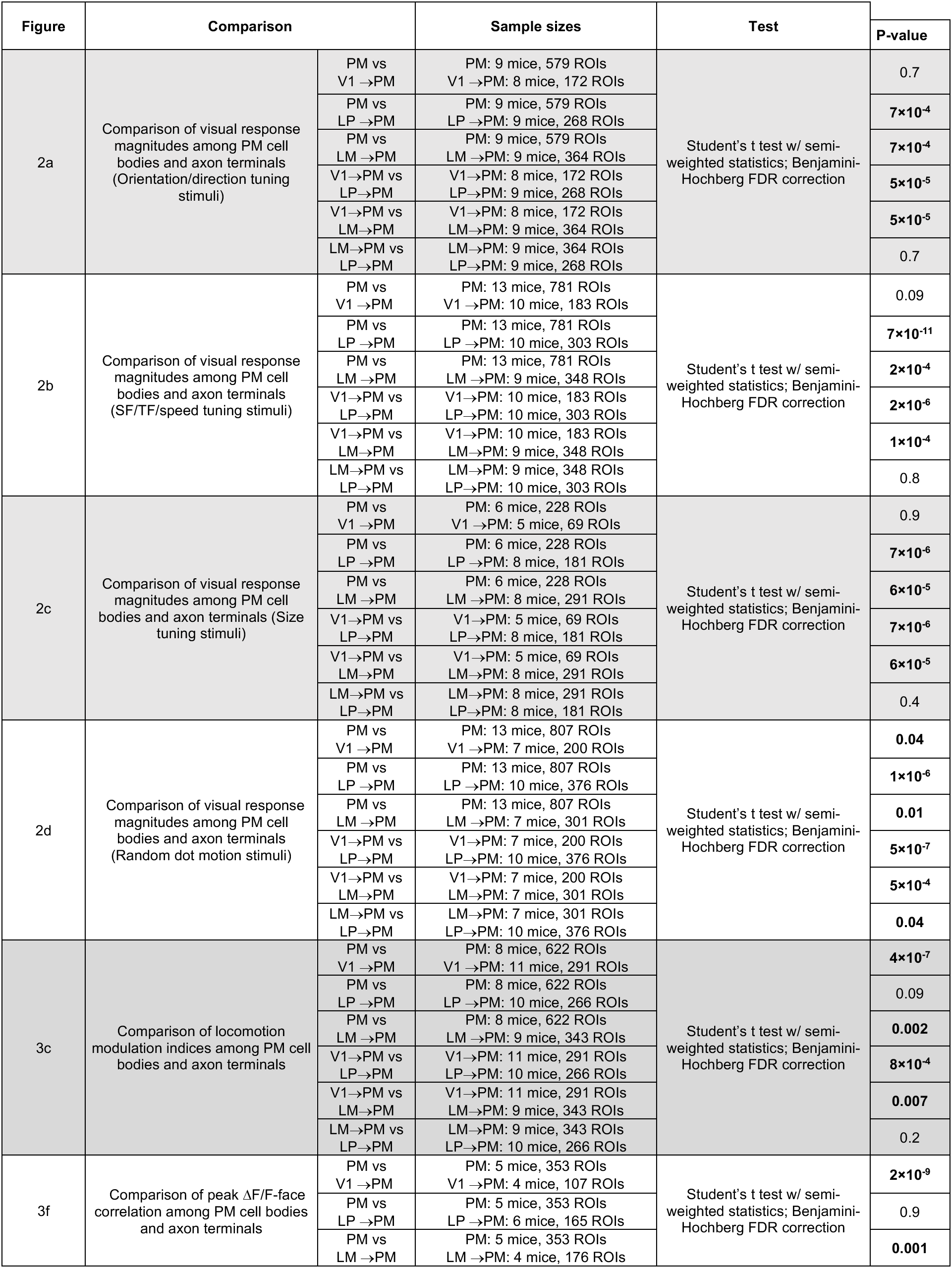
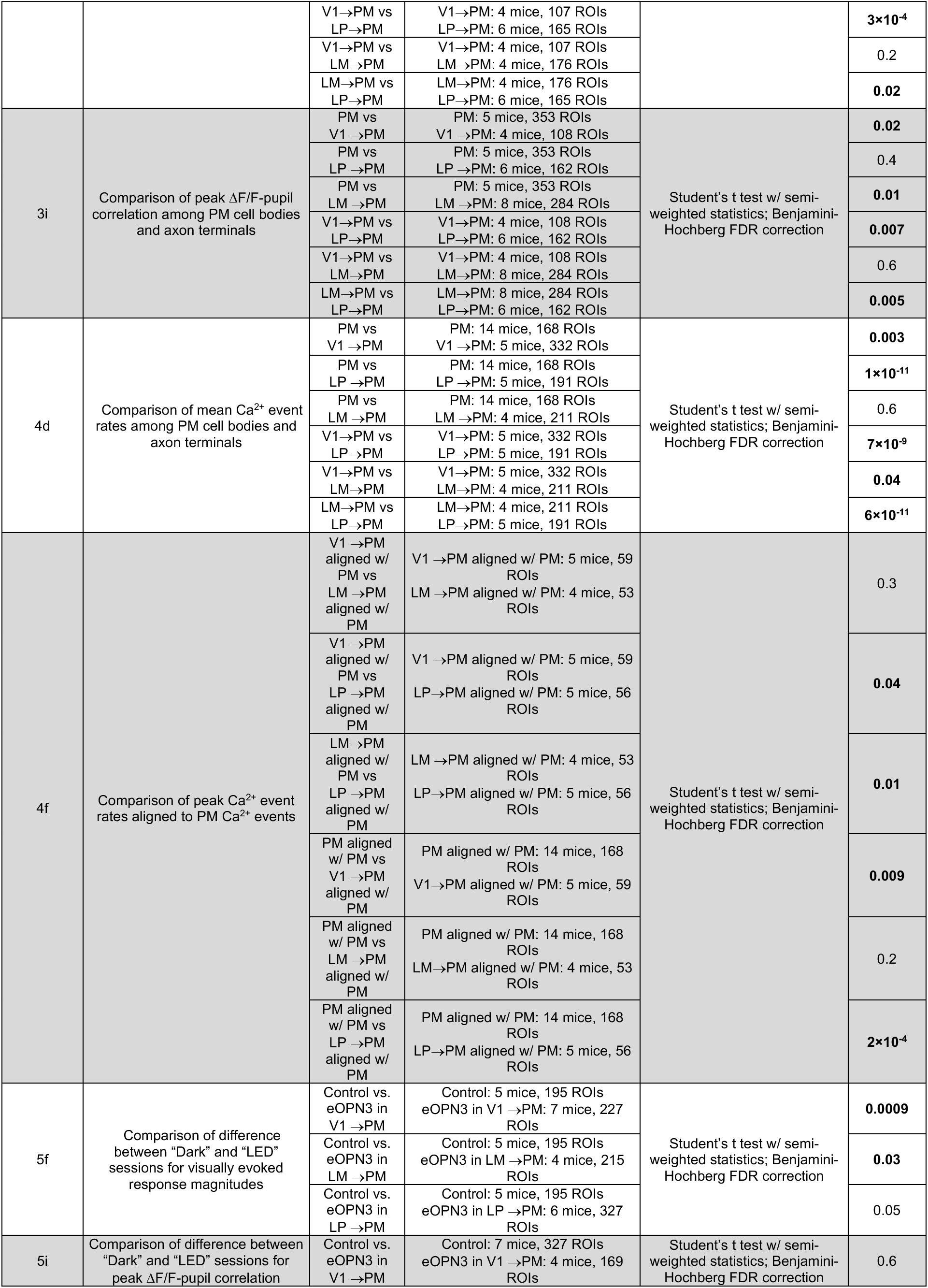
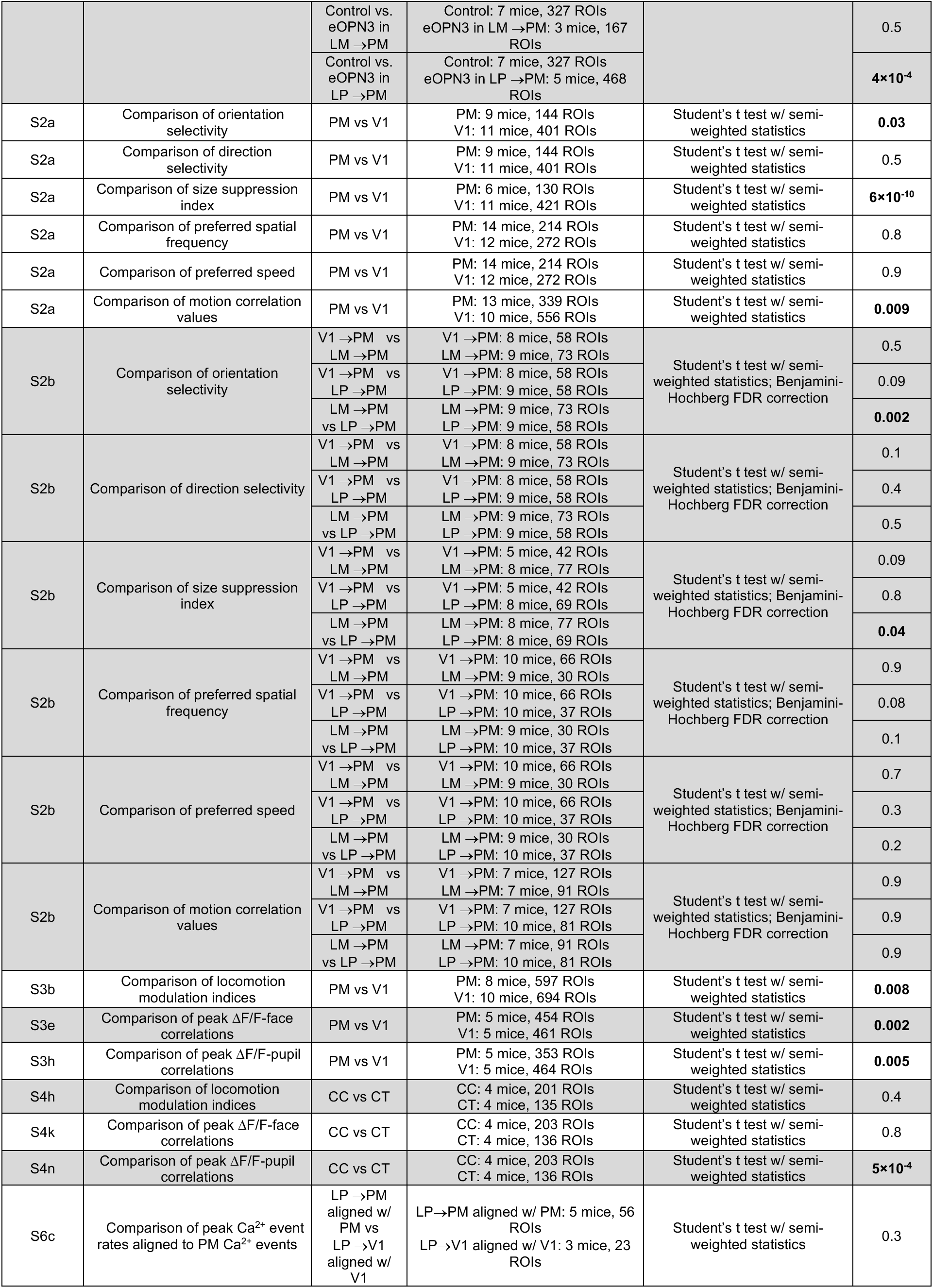
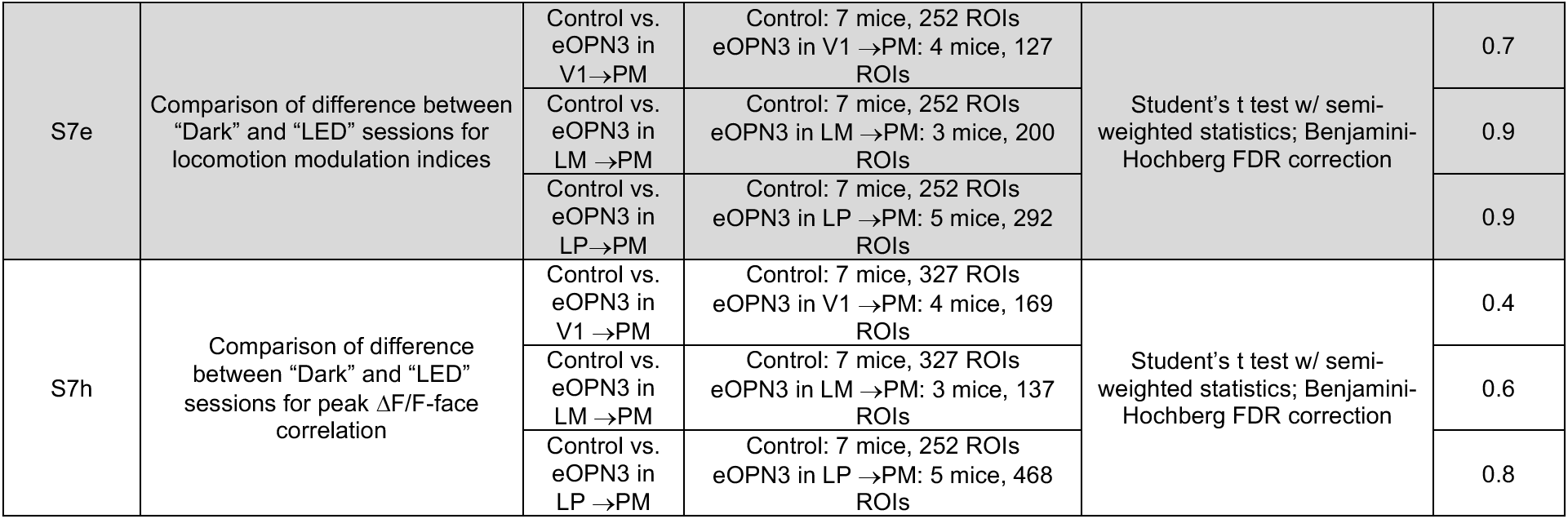
Summary of all statistical analyses.

## Notes

### Competing Interest Statement

The authors have declared no competing interest.

## REFERENCES

Aarts E, Verhage M, Veenvliet JV, Dolan CV, van der Sluis S (2014) A solution to dependency: using multilevel analysis to accommodate nested data. Nat. Neurosci. 17: 491–496.

Andermann ML, Kerlin AM, Roumis DK, Glickfeld LL, Reid RC (2011) Functional specialization of mouse higher visual cortical areas. Neuron. 72:1025–1039.

Bayer L, Serafin M, Eggermann E, Saint-Mleux B, Machard D, Jones BE, Mühlethaler M (2004) Exclusive postsynaptic action of hypocretin-orexin on sublayer 6b cortical neurons. J. Neurosci. 24: 6760–6764.

Beaulieu-Laroche L, Toloza EHS, Brown NJ, Harnett MT (2019) Widespread and highly correlated somato-dendritic activity in cortical layer 5 neurons. Neuron 103: 235–241.

Beltramo R, Scanziani M (2019) A collicular visual cortex: neocortical space for an ancient midbrain visual structure. Science 363: 64–69.

Benisty H, Barson A, Moberly AH, Lohani S, Coifman RR, Mishne G, Crair MC, Cardin JA, Higley MJ (2023) Rapid fluctuations in functional connectivity of cortical networks encode spontaneous behavior. bioRxiv.

Bennett C, Gale SD, Garrett ME, Newton ML, Callaway EM, Murphy GJ, Olsen SR (2019) Higher-order thalamic circuits channel parallel streams of visual information in mice. Neuron 102: 477–492.

Blot A, Roth MM, Gasler I, Javadzadeh M, Imhof F, Hofer SB (2021) Visual intracortical and transthalamic pathways carry distinct information to cortical areas. Neuron 109: 1996–2008.

Bolkor H, Frère SGA, Eyre MD, Slézia A, Ulbert I, Lüthi A, Acsády L (2005) Selective GABAergic control of higher-order thalamic relays. Neuron 45: 929–940.

Brainard DH (1997) The psychophysics toolbox. Spat. Vis. 10: 433–436.

Chen TW, Wardill TJ, Sun Y, Pulver SR, Renninger SL, Baohan A, Schreiter ER, Kerr RA, Orger MB, Jayaraman V, Looger LL, Svoboda K, Kim DS (2013) Ultrasensitive fluorescent proteins for imaging neuronal activity. Nature 499: 295–300.

Chung Y, Rabe-Hesketh S, Choi IH (2013) Avoiding zero between-study variance estimates in random-effects meta-analysis. Stat. Med. 32: 4071–4089.

Collins L, Francis J, Emanuel B, McCormick DA (2023) Cholinergic and noradrenergic axonal activity contains a behavioral-state signal that is coordinated across the dorsal cortex. Elife 12: e81826.

Crandall SR, Cruikshank SJ, Connors BW (2015) A corticothalamic switch: controlling the thalamus with dynamic synapses. Neuron 86: 768–782.

D’Souza RD, Meier AM, Bista P, Wang Q, Burkhalter A (2016) Recruitment of inhibition and excitation across mouse visual cortex depends on the hierarchy of interconnecting areas. Elife 5: e19332.

D’Souza RD, Wang Q, Ji, W, Meier AM, Kennedy H, Knoblauch K, Burkhalter A (2022) Hierarchical and nonhierarchical features of the mouse visual cortical network. Nat. Commun. 13: 503.

Dubbs A, Guevara J, Yuste R (2016) moco: fast motion correction for calcium imaging. Front. Neuroinform. 10: 6.

El-Shamayleh Y, Kumbhani RD, Dhruv NT, Movshon JA (2013) Visual response properties of V1 neurons projecting to V2 in macaque. J. Neurosci. 33: 16594–16605.

Felleman DJ, Van Essen DC (1991) Distributed hierarchical processing in the primate cerebral cortex. Cereb. Cortex 1: 1–47.

Ferguson KA, Salameh J, Alba C, Selwyn H, Barnes C, Lohani S, Cardin JA (2023) VIP interneurons regulate cortical size tuning and visual perception. bioRxiv.

Fiebelkorn IC, Pinsk MA, Kastner S (2019) The mediodorsal pulvinar coordinates the macaque fronto-parietal network during rhythmic spatial attention. Nat. Commun. 10: 215.

Friedrich J, Zhou P, Paninski L (2017) Fast online deconvolution of calcium imaging data. PLoS Comput. Biol. 13: e1005423.

Fu Y, Tucciarone JM, Espinosa JS, Sheng N, Darcy DP, Nicoll RA, Huang ZJ, Stryker MP (2014) A cortical circuit for gain control by behavioral state. Cell 156: 1139 –1152.

Gămănuţ R, Kennedy H, Toroczkai Z, Ercsey-Ravasz M, Van Essen DC, Knoblauch K, Burkhalter A (2018) The mouse cortical connectome, characterized by an ultra-dense cortical graph, maintains specificity by distinct connectivity profiles. Neuron 97: 698–715.

Glickfeld LL, Andermann ML, Bonin V, Reid RC (2013) Cortico-cortical projections in mouse visual cortex are functionally target specific. Nat. Neurosci. 16: 219–226.

Glickfeld LL, Olsen SR (2017) Higher-order areas of the mouse visual cortex. Annu. Rev. Vis. Sci. 3: 251–273.

Goltstein PM, Reinert S, Bonhoeffer T, Hübener M. (2021) Mouse visual cortex areas represent perceptual and semantic features of learned visual categories. Nat. Neurosci. 24:1441–51.

Groh A, de Kock CP, Wimmer VC, Sakmann B, Kuner T (2008) Driver or coincidence detector: modal switch of a corticothalamic giant synapse controlled by spontaneous activity and short-term depression. J. Neurosci. 28: 9652–9663.

Grujic N, Tesmer A, Bracey E, Peleg-Raibstein D, Burdakov D (2023) Control and coding of pupil size by hypothalamic orexin neurons. Nat. Neurosci. 26: 1160–1164.

Haider B, McCormick DA (2009) Rapid neocortical dynamics: cellular and network mechanisms. Neuron 62: 171–189.

Harris KD, Shepherd GMG (2015) The neocortical circuit: themes and variations. Nat. Neurosci. 18: 170–181.

Hoerder-Suabedissen A, Hayashi S, Upton L, Nolan Z, Casas-Torremocha D, Grant E, Viswanathan S, Kanold PO, Clasca F, Kim Y, Molnár Z (2018) Subset of cortical layer 6b neurons selectively innervates higher order thalamic nuclei in mice. Cereb. Cortex 28: 1882–1897.

Junyent F, Kremer EJ (2015) CAV-2—why a canine virus is a neurobiologist’s best friend. Curr. Opin. Pharmacol. 24: 86–93.

Kaas JH, Lyon DC (2007) Pulvinar contributions to the dorsal and ventral streams of visual processing in primates. Brain Res. Rev. 55: 285–296.

Kohn A, Jasper AI, Semedo JD, Gokcen E, Machens CK, Yu BM (2020) Principles of corticocortical communication: proposed schemes and design considerations. Trends Neurosci. 43: 725–737.

LaTerra D, Bjerre AS, Rosier M, Masuda R, Ryan TJ, Palmer LM (2022) The role of higher-order thalamus during learning and correct performance in goal-directed behavior. Elife 11: e77177.

Li J, Guido W, Bickford ME (2003) Two distinct types of corticothalamic EPSPs and their contribution to short-term synaptic plasticity. J. Neurophysiol. 90: 3429–3440.

Li JY, Hass CA, Matthews I, Kristl AC, Glickfeld LL (2021) Distinct recruitment of feedforward and recurrent pathways across higher-order areas of mouse visual cortex. Curr. Biol. 31: 5024–5036.

Lohani S, Moberly AH, Benisty H, Landa B, Jing M, Li Y, Higley MJ, Cardin JA (2022) Spatiotemporally heterogeneous coordination of cholinergic and neocortical activity. Nat. Neurosci. 25: 1706–1713.

Luo L, Callaway EM, Svoboda K (2018) Genetic dissection of neural circuits: a decade of progress. Neuron 98: 256–281.

Lur G, Vinck MA, Tang L, Cardin JA, Higley MJ (2016) Projection-specific visual feature encoding by layer 5 cortical subnetworks. Cell Rep. 14: 2538–2545.

Orban GA (2008) Higher order visual processing in macaque extrastriate cortex. Physiol. Rev. 88: 59–89.

Mahn M, Saraf-Sinik I, Patil P, Pulin, M, Bitton E, Karalis N, Bruentgens F, Palgi S, Gat A, Dine J, Wietek J, Davidi I, Levy R, Litvin A, Zhou F, Sauter K, Soba P, Schmitz D, Lüthi A, Rost BR, Wiegert JS, Yizhar O (2021) Efficient optogenetic silencing of neurotransmitter release with a mosquito rhodopsin. Neuron 109: 1621–1635.

Martinez-Garcia RI, Voelcker B, Zaltsman JB, Patrick SL, Stevens TR, Connors BW, Cruikshank SJ (2020) Two dynamically distinct circuits drive inhibition in the sensory thalamus. Nature 583: 813–818.

Mazurek M, Kager M, Van Hooser SD (2014) Robust quantification of orientation selectivity and direction selectivity. Front. Neural Circuits 8: 92

McCormick DA (1992) Neurotransmitter actions in the thalamus and cerebral cortex and their role in neuromodulation of thalamocortical activity. Prog. Neurobiol. 39: 337–388.

McGinley MJ, Vinck M, Reimer J, Batista-Brito R, Zagha E, Cadwell CR, Tolias AS, Cardin JA, McCormick DA (2015) Waking state: rapid variations modulate neural and behavioral responses. Neuron 87:1143–1161.

Merabet L, Desautels A, Minville K, Casanova C (1998) Motion integration in a thalamic visual nucleus. Nature 396: 265–268.

Miller-Hansen AJ, Sherman SM (2022) Conserved patterns of functional organization between cortex and thalamus in mice. Proc. Natl. Acad. Sci. USA 119: e2201481119.

Molnár B, Sere P, Bordé S, Koós K, Zsigri N, Horváth P, Lőrincz ML. (2021) Cell type-specific arousal-dependent modulation of thalamic activity in the lateral geniculate nucleus. Cerebral Cortex Communications. 2:tgab020.

Movshon JA, Newsome WT (1996) Visual response properties of striate cortical neurons projecting to area MT in macaque monkeys. J. Neurosci. 16: 7733–7741.

Murgas, KA, Wilson AM, Michael V, Glickfeld LL (2020) Unique spatial integration in mouse primary visual cortex and higher visual areas. J. Neurosci. 40: 1862–1873

Musall S, Kaufman MT, Juavinett AL, Gluf S, Churchland AK (2019) Single-trial neural dynamics are dominated by richly varied movements. Nat. Neurosci. 22: 1677–1686.

Musall S, Sun XR, Mohan H, Gluf S, Li S, Drewes R, Cravo E, Lenzi I, Yin C, Kampa BM, Churchland AK (2023) Pyramidal cell types drive functionally distinct cortical activity patterns during decision-making. Nat. Neurosci. 26:495–505.

Neske GT, Nestvogel D, Steffan PJ, McCormick DA (2019) Distinct waking states for strong evoked responses in primary visual cortex and optimal visual detection performance. J. Neurosci. 39: 10044–10059.

Nestvogel DB, McCormick DA (2022) Visual thalamocortical mechanisms of waking state-dependent activity and alpha oscillations. Neuron 110: 120–138.

Niell CM, Stryker MP (2008) Highly selective receptive fields in mouse visual cortex. J. Neurosci. 28: 7520–7536.

Niell CM, Stryker MP (2010) Modulation of visual responses by behavioral state in mouse visual cortex. Neuron 65:472–479.

Paxinos G, Franklin KBJ (2008) The mouse brain in stereotaxic coordinates. Ed. 3. (Academic Press).

Peters AJ, Lee J, Hedrick NG, O’Neil K, Komiyama T (2017) Reorganization of corticospinal output during motor learning. Nat. Neurosci. 20: 1133–1141.

Petreanu L, Mao T, Sternson SM, Svoboda K (2009) The subcellular organization of neocortical excitatory connections. Nature 457: 1142–1145.

Petty GH, Kinnischtzke AM, Hong YK, Bruno RM (2021) Effects of arousal and movement on secondary somatosensory and visual thalamus. Elife 10: e67611.

Polack PO, Friedman J, Golshani P (2013) Cellular mechanisms of brain state-dependent gain modulation in visual cortex. Nat. Neurosci. 16:1331–1339.

Poulet JA, Fernandez LMJ, Crochet S, Petersen CCH (2012) Thalamic control of cortical states. Nat. Neurosci. 15: 370–372.

Priebe NJ, Lisberger SG, Movshon JA (2006) Tuning for spatiotemporal frequency and speed in directionally selective neurons of macaque striate cortex. J. Neurosci. 26: 2941–2950.

Reichova I, Sherman SM (2004) Somatosensory corticothalamic projections: distinguishing drivers from modulators. J. Neurophysiol. 92: 2185–2195.

Reimer J, Froudarakis E, Cadwell CR, Yatsenko D, Denfield GH, Tolias AS (2014) Pupil fluctuations track fast switching of cortical states during quiet wakefulness. Neuron 84:355–362.

Reimer J, McGinley MJ, Liu Y, Rodenkirch C, Wang Q, McCormick DA, Tolias AS (2016) Pupil fluctuations track rapid changes in adrenergic and cholinergic activity in cortex. Nat. Commun. 7: 13289.

Reinhold K, Resulaj A, Scanziani M. (2023) Brain state-dependent modulation of thalamic visual processing by cortico-thalamic feedback. J. Neurosci. 43:1540–54.

Riesenhuber M, Poggio T (1999) Hierarchical models of object recognition in cortex. Nat. Neurosci. 2:1019–1025.

Roth MM, Dahmen JC, Muir DR, Imhof F, Martini FJ, Hofer SB (2016) Thalamic nuclei convey diverse contextual information to layer 1 of visual cortex. Nat. Neurosci. 19: 299–307.

Saalmann YB, Pinsk MA, Wang L, Li X, Kastner S (2012) The pulvinar regulates information transmission between cortical areas based on attention demands. Science 337: 753–756.

Salkoff DB, Zagha E, McCarthy E, McCormick DA (2020) Movement and performance explain widespread cortical activity in a visual detection task. Cereb. Cortex 30: 421–437.

Schmitt LI, Wimmer RD, Nakajima M, Happ M, Mofakham S, Halassa MM (2017) Thalamic amplification of cortical connectivity sustains attentional control. Nature 545: 219–223.

Sherman SM, Guillery RW (2011) Distinct functions for direct and transthalamic corticocortical connections. J. Neurophysiol. 106: 1068–1077.

Shipp S (2003) The functional logic of cortico-pulvinar connections. Philos. Trans. R. Soc. Lond. B Biol. Sci. 358: 1605–1624.

Siegle JH, Jia X, Durand S, Gale S, Bennett C, et al. (2021) Survey of spiking in the mouse visual system reveals functional hierarchy. Nature 592: 86–92.

Sit KK, Goard MJ (2020) Distributed and retinotopically asymmetric processing of coherent motion in mouse visual cortex. Nat. Commun. 11: 3565.

Spacek MA, Crombie D, Bauer Y, Born G, Liu X, Katzner S, Busse L. (2022) Robust effects of corticothalamic feedback and behavioral state on movie responses in mouse dLGN. Elife. 11:e70469.

Stringer C, Pachitariu M, Steinmetz B, Reddy CB, Carandini M, Harris KD (2019) Spontaneous behaviors drive multidimensional, brainwide activity. Science 364: 255.

Tang L, Higley MJ (2020) Layer 5 circuits in V1 differentially control visuomotor behavior. Neuron 105: 346–354.

Theyel BB, Llano DA, Sherman SM (2010) The corticothalamocortical circuit drives higher-order cortex in the mouse. Nat. Neurosci. 13: 84–88.

Ungerleider LG, Mishkin M (1982) Two cortical visual systems. In Analysis of Visual Behavior, DJ Ingle, MA Goodale, RJW Mansfield, eds. (MIT Press), pp. 549–586.

Van Essen DC (2005) Corticocortical and thalamocortical information flow in the primate visual system. Prog. Brain Res. 149: 173–185.

Varela C (2014) Thalamic neuromodulation and its implications for executive networks. Front. Neural Circuits 8: 69.

Varela C, Sherman SM (2007) Differences in response to muscarinic agonists between first and higher-order thalamic relays. J. Neurophysiol. 98: 3538–3547.

Varela C, Sherman SM (2009) Differences in response to serotonergic activation between first and higher-order thalamic nuclei. Cereb. Cortex 19: 1776–1786.

Vinck M, Batista-Brito R, Knoblich U, Cardin JA (2015) Arousal and locomotion make distinct contributions to cortical activity patterns and visual encoding. Neuron 86: 740–754.

Wang Q, Burkhalter A (2007) Area map of mouse visual cortex. J. Comp. Neurol. 502: 339357.

Wang Q, Sporns O, Burkhalter A (2012) Network analysis of corticocortical connections reveals ventral and dorsal processing streams in mouse visual cortex. J. Neurosci. 32: 4386–4399.

Wang Q, Gao E, Burkhalter A (2011) Gateways of ventral and dorsal streams in mouse visual cortex. J. Neurosci. 31: 1905–1918.

Xu X, Holmes TC, Luo MH, Beier KT, Horwitz GD, Zhao F, Zeng W, Hui M, Semler BL, Sandri-Goldin RM (2020) Viral vectors for neural circuit mapping and recent advances in trans-synaptic anterograde tracers. Neuron 107: 1029–1047.

Zagha E, Casale AE, Sachdev RNS, McGinley MJ, McCormick DA (2013) Motor cortex feedback influences sensory processing by modulating network state. Neuron 79: 567–578.

Zhou H, Schafer RJ, Desimone R (2016) Pulvinar-cortex interactions in vision and attention. Neuron 89: 209–220.

Zhuang J, Ng L, Williams D, Valley M, Li Y, Garrett M, Waters J (2017) An extended retinotopic map of mouse cortex. Elife 6: e18372.

Zingg B, Hintiryan H, Gou L, Song MY, Bay M, Bienkowski MS, Foster NN, Yamashita S, Bowman I, Toga AW, Dong HW (2014) Neural networks of the mouse neocortex. Cell 156: 1096–1111.

